# T cells Instruct Immune Checkpoint Inhibitor Therapy Resistance in Tumors Responsive to IL-1 and TNFα Inflammation

**DOI:** 10.1101/2022.09.20.508732

**Authors:** Nam Woo Cho, Sophia M. Guldberg, Barzin Y. Nabet, Jie Zeng Yu, Eun Ji Kim, Kamir J. Hiam-Galvez, Jacqueline L. Yee, Rachel DeBarge, Iliana Tenvooren, Naa Asheley Ashitey, Filipa Lynce, Deborah A. Dillon, Jennifer M. Rosenbluth, Matthew H. Spitzer

**Author notes:** **Corresponding authors:** Matthew H. Spitzer UCSF Box 0552, 513 Parnassus Ave., San Francisco, CA 94143, USA Nam Woo Cho 2340 Sutter St., San Francisco, CA 94115, USA.

## Abstract

Resistance to immune checkpoint inhibitors (ICIs) is common, even in tumors with T cell infiltration. We thus investigated consequences of ICI-induced T cell infiltration in the microenvironment of resistant tumors. T cells increased in ICI-resistant tumors following treatment as did neutrophils, in contrast to ICI-responsive tumors. Resistant tumors were distinguished by high expression of IL-1 Receptor 1 (IL1R1), enabling a synergistic response to IL-1 and TNFα to induce G-CSF, CXCL1, and CXCL2 via NF-κB signaling, supporting neutrophils. Perturbation of this inflammatory resistance circuit sensitized tumors to ICIs. Paradoxically, T cells drove this resistance circuit via TNFα both *in vitro* and *in vivo*. Evidence of this inflammatory resistance circuit and its impact also translated to human cancers. These data support a novel mechanism of ICI resistance, wherein treatment-induced T cell activity can drive resistance in tumors responsive to IL-1 and TNFα, with important therapeutic implications.

**Statement of Significance:** Although T cell-infiltrated cancers are frequently resistant to immune checkpoint inhibitor therapies, mechanisms of resistance beyond T cell exhaustion remain unclear. Here, we reveal the functional significance of tumor- infiltrating T cells in resistant tumors, which surprisingly instruct immunosuppressive inflammation in mouse and human cancers responsive to IL-1 and TNFα.

## Introduction

Immune checkpoint inhibitors (ICIs) are integral to the armamentarium against cancer. However, these therapies are not effective in many patients, and an important mission is to better define principles determining treatment response versus failure^1^. Among core immunologic programs governing the response to ICIs is the infiltration of T cells in the tumor microenvironment (TME)^2–5^, where “inflamed” or “hot” tumors, such as those with microsatellite instability-high (MSI-H) status, exhibit higher response rates to further immune activating therapies^6–10^. Nevertheless, treatment resistance to ICIs occurs commonly even in T cell-infiltrated tumors^11,12^, but there is a limited understanding of how such T cell-infiltrated tumors can escape T cell-activating therapies. For example, MSI-H tumors demonstrate increased tumor neoantigen load and T cell infiltration, yet not all respond to ICIs^9–11,13–18^. We reasoned that the development of T cell infiltrated tumor models would enable study of T cell states and their interactions with the TME during ICI resistance despite T cell inflammation.

Inflammation is a broad term encompassing multiple processes that can promote or prevent tumor progression^19^. For instance, inflammation can be driven by the transcription factor nuclear factor-κB (NF-κB), which is activated in multiple cell types in the TME by various stimuli, including the cytokines TNFα and IL- 1^20,21^. This circuit can drive diverse tumor promoting mechanisms including immunosuppression^22–26^. Among many known roles, NF-κB signaling can foster recruitment and activation of immunosuppressive myeloid cells to suppress T cell cytotoxicity^27–31^. Accordingly, its inhibition in the TME can lead to improved immunologic control of tumors^31–36^. Of note, T cells and NF-κB-mediated immunosuppressive inflammation can co-exist in the TME, yet their relationship is incompletely defined. Much of the recent research efforts have focused on the biology of tumor infiltrating lymphocytes (TILs), which lose stemness and differentiate toward exhaustion as a result of chronic antigen stimulation and immunosuppressive signals, including those provided by myeloid cell subsets^37,38.^. However, it is currently unclear whether and how conventional CD4 or CD8 T cells themselves regulate inflammation in the TME, and whether the interplay between these factors shapes the immunologic setpoint to affect responsiveness to immunotherapy.

Here, we use T cell-infiltrated mouse syngeneic tumor models of ICI response and resistance to discover that T cells can drive a TNFα and IL-1-mediated inflammatory response from IL1R1-expressing tumor cells in the setting of ICI resistance. TNFα and IL-1 synergistically upregulate NF-κB-transcribed cytokines including G-CSF, CXCL1, and CXCL2, resulting in recruitment of immunosuppressive neutrophils to the TME. Importantly, we show that this inflammatory resistance circuit requires T cells. Activated tumor-infiltrating T cells were sufficient to increase G-CSF production by tumor cells *in vitro* in a TNFα-dependent manner. *In vivo,* ICIs increased infiltration of TNFα-producing T cells. Breaking this inflammatory circuit via pharmacologic or genetic perturbation of IL1R1 or TNFα, including tumor cell-specific deletion of IL1R1 or T cell-specific deletion of TNFα, sensitized resistant tumors to ICIs. Finally, analyses of human datasets corroborate these cellular and mechanistic elements in patients treated with ICIs, revealing correlation of ICI-induced infiltration of *TNF+* T cells and NF-κB activity in IL1R1-expressing tumors, as well as the adverse impact of the inflammatory resistance axis on ICI treatment outcome. These findings refine the current paradigm of T cell responses to ICI therapy with significant clinical implications, revealing an opportunity to dissociate the tumor-promoting aspects of T cell activity from their tumoricidal functions to improve therapeutic benefit.

## Results

### T Cell-Infiltrated Mouse Syngeneic Tumors Demonstrate Variable Responses to ICIs

Tumors lacking T cell infiltration are generally resistant to ICIs. While the opposite may be expected for tumors with high T cell infiltration, a significant fraction nevertheless demonstrates treatment resistance^11,12^. To understand mechanisms of ICI response and resistance in T cell-infiltrated tumors, we generated mouse syngeneic tumor cell lines lacking the DNA mismatch repair protein, MSH2, to create microsatellite instability-high (MSI- H) cells. While the response rates to ICIs are generally favorable, treatment resistance is nevertheless frequently seen in patients with response rates ranging from 30-50% in pre-treated settings^11,17,18^, and in mouse MSI-H tumor models^39,40^. Thus, we utilized this opportunity to study ICI response and resistance in the context of T cell infiltration. *Msh2* gene deletion by CRISPR-Cas9 ribonucleoprotein electroporation^41^ in the polyclonal cell pool almost completely abolished MSH2 protein expression, eliminating the requirement for single cell cloning and allowing preservation of any parental tumor heterogeneity, which has been shown to impact immunogenicity and ICI sensitivity^42^ (Fig. 1A and B; Supplementary Fig. S1A; Supplementary Table S1). Cell lines generated in this manner accumulated microsatellite instability assessed by PCR^9,43^ over 4 months, or 60 passages, of continuous culture (Fig. 1C; Supplementary Fig. S1B; Supplementary Table S2). Compared to control microsatellite-stable (MSS) cell lines generated using nontargeting guide RNA and cultured for the same duration, the MSI-H cell lines accumulated single nucleotide polymorphisms and indels when quantified by whole exome sequencing (Fig. 1D). These data are consistent with the level of MSI reported in published mouse studies^9,10^. Each tumor cell line transcribed neoantigens with a similar distribution and heterogeneity of predicted antigenicity and allele frequency, calculated using MuPeXI with inputs including neopeptide sequences, predicted MHC binding affinities and gene expression^44^ (Fig. 1E; Supplementary Fig. S1C). Loss of MHC class I antigen presentation or responsiveness to IFNγ are established, albeit somewhat rare, mechanisms of ICI resistance^45^. However, each model cell line also maintained the ability to induce MHC-I expression upon IFNγ stimulation (Supplementary Fig. S1D).

**Figure 1:**
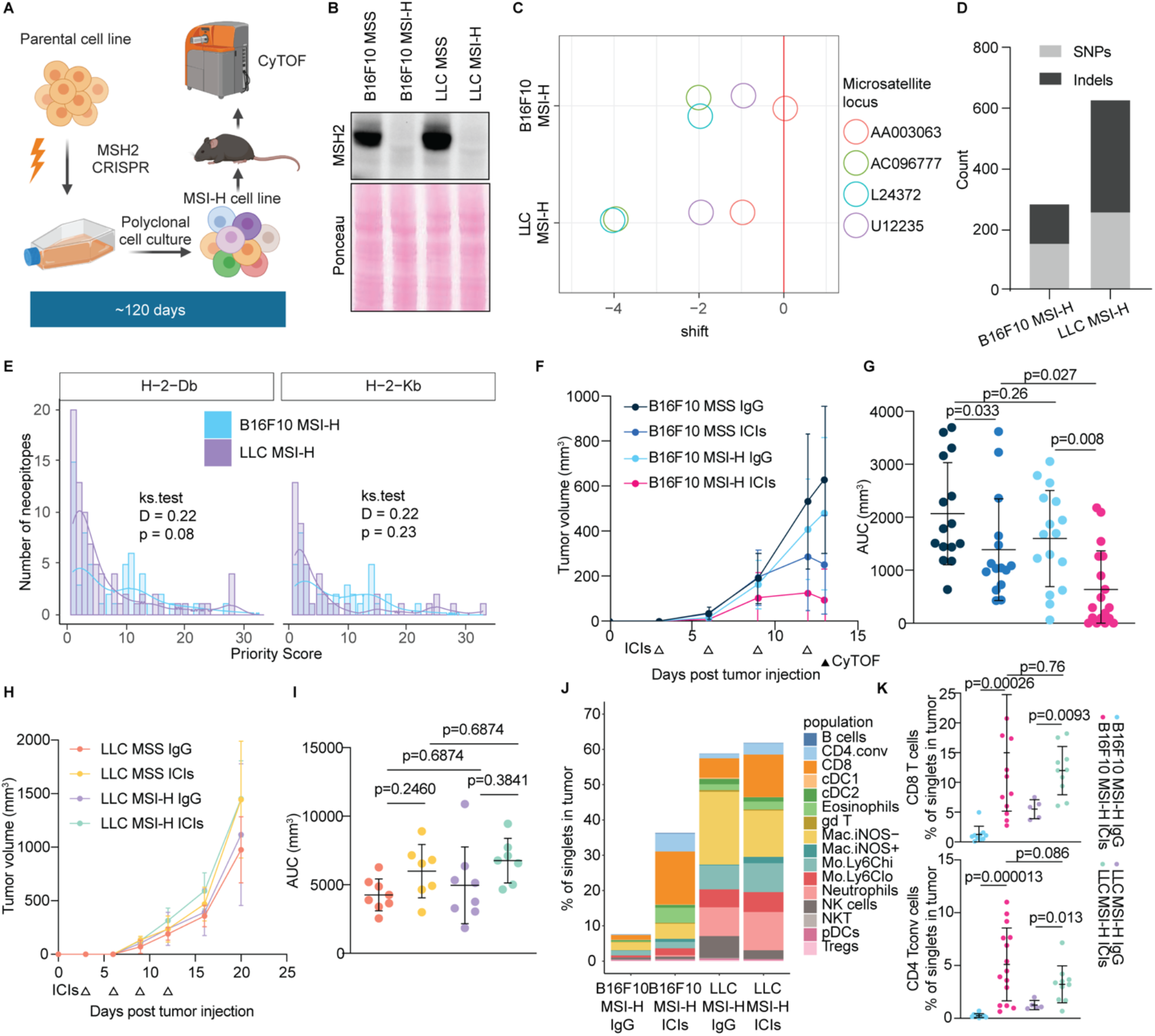
T cell-infiltrated tumors demonstrate variable responses to ICIs. **A,** Outline for generation and experimentation of MSI-H syngeneic tumors. **B,** Representative western blot of whole cell lysates of indicated cell lines. 3 independent experiments were performed. **C,** Targeted microsatellite PCR assay comparing amplicon size at indicated loci for MSI-H vs. MSS cell line. Representative of two independent experiments. **D,** Whole exome sequencing quantification of single nucleotide polymorphisms (SNPs) and insertion-deletion mutations (indels) for indicated MSI-H cell lines compared to the MSS lines. **E,** MuPeXI neoepitope affinity scores for indicated cell lines and HLA class. D statistic and p values are calculated using a two-sided Kolmogorov-Smirnov test. **F and G,** Quantification of mean volumes and area under curves (AUC) for B16F10 subcutaneous tumors treated with or without anti-PD1 and anti-CTLA4 antibodies (ICIs) at the indicated timepoints (white arrowhead). Black arrowhead denotes timepoint for sample collection for CyTOF analysis. n=15-18 mice per condition from two independent experiments. Two-tailed adjusted p value by Mann-Whitney tests. **H and I,** Analysis as in **(F)** and **(G)** for subcutaneous LLC tumors. n=7-8 mice per condition from one experiment. Two-tailed adjusted p values by Welch’s t tests. **J,** Immune subsets in tumor were defined by manual gating of CyTOF data from B16F10 subcutaneous or LLC MFP tumors harvested at day 13 post implantation. Conv=conventional; Mac=macrophages; Mo=monocytes. Bar height for each population represents the mean of biological replicates (n=5-15 mice) from three independent experiments. **K,** Quantification of CD8 (top panel) and CD4 Tconv (bottom panel) infiltration from **(J)**. Two-tailed adjusted p value by Mann-Whitney tests. For all applicable panels, error bars represent mean +/- SD.

Notably, when implanted into syngeneic C57BL/6J mice, the tumor models demonstrated contrasting responses to combination ICI treatment of anti-PD-1 and anti-CTLA4, paralleling the observed ICI response rates for human MSI-H cancers^11,15^. The B16F10 melanoma subcutaneous tumors responded to ICIs, where MSI-H tumors were more sensitive to ICIs compared to MSS tumors, although *in vitro* cell doubling times were not different between MSS and MSI-H cells (Fig. 1F and G; Supplementary Fig. S1E and F). The EO771 breast MSS and MSI-H tumors, implanted into the mammary fat pad, also responded to ICIs, with the greatest response seen in MSI-H tumors (Supplementary Fig. S1G-I). However, resistance to ICIs was notable in the MSS or MSI-H LLC tumors in either the subcutaneous space or the mammary fat pad, sites of implantation that result in similar tumor immune composition (Fig. 1H and I; Supplementary Fig. S1J-N).

To investigate mechanisms responsible for the divergent ICI responses in these models, we used mass cytometry by time-of-flight (CyTOF), analyzing the immune infiltrate from tumor samples on day 13 post- implantation, where tumor sizes are similar across the models. Identification of major immune cell types in the TME revealed that ICI treatment induced T cell infiltration in both B16F10 and LLC MSI-H tumors (Fig. 1J and K, Supplementary Fig. S2), suggesting that the ICI resistance in the LLC MSI-H model is unlikely to be attributable solely to insufficient numbers of TILs. ICI resistance in T cell-infiltrated tumors was not exclusive to the MSI-H context or the C57BL/6J genotype. EMT6 breast tumors in the mammary fat pad of BALB/cJ mice were also resistant to combination ICIs, despite increased CD8 T cell infiltration following treatment (Supplementary Fig. S3A-C).

### ICI-Resistant Tumors Acquire Unfavorable T cell Activation States After Treatment

To define phenotypic differences in TILs in ICI-responsive vs. resistant tumors, we further analyzed antigen-experienced CD44+ CD8 and conventional CD4 T cells (CD4 Tconv) by Uniform Manifold Approximation and Projection (UMAP)^46^, and by differential abundance (DA) analysis^47^. This approach enabled identification of cell subsets enriched in each condition and definition of their differentiation and activation states. We focused our analysis to understand the differences between the ICI-treated B16F10 MSI-H and LLC MSI-H tumors. Analysis of day 13 tumor-infiltrating CD8 T cells showed differentially-abundant (DA) cells for each condition (Fig. 2A and B) with distinct marker expression (Fig. 2C). To further quantify the phenotypic differences, DA cells were subsequently grouped by unsupervised clustering. Clusters 4, 5 and 6, which were enriched in B16F10 MSI-H tumors, were notably higher in expression of markers characteristic of activated effector T cells, including Ki67, granzyme B (GrB), CD39, CD38, and PD1. In contrast, cluster 2, which was enriched in LLC tumors, expressed comparatively lower expression of these markers (Fig. 2D and E). LLC tumors were also enriched in cluster 3 cells, which expressed TCF1 but were PD1-, resembling memory precursor-like CD8 T cells that have been shown to demonstrate more polyfunctionality, including TNFα production^48^.

**Figure. 2:**
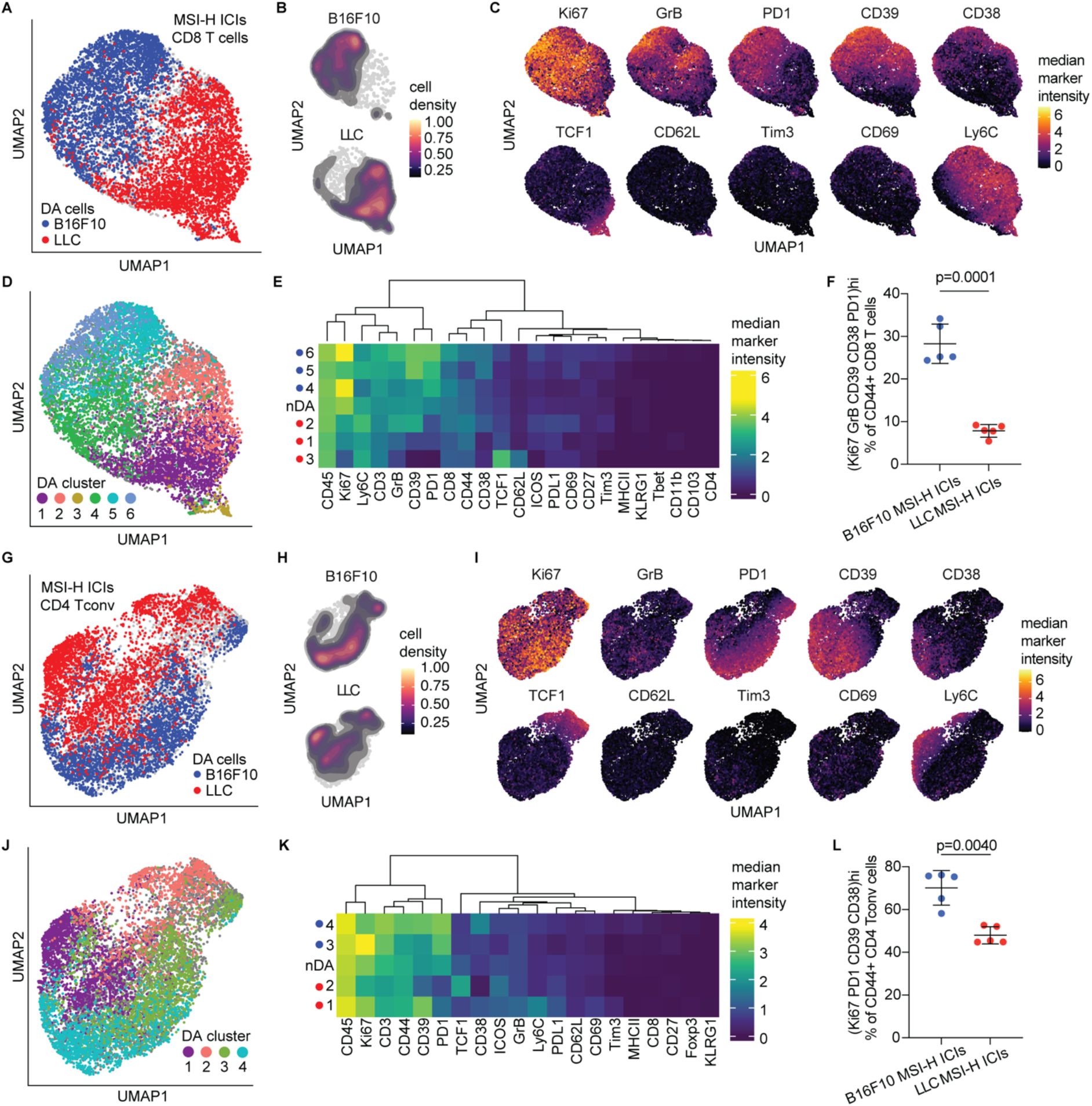
Distinct effector marker expression in T cells in ICI-responsive vs. ICI-resistant tumors. **A,** Differential abundance (DA) analysis of 10,000 CD44+ CD8 T cells from ICI-treated B16F10 MSI-H and LLC MSI-H tumors harvested at day 13. CyTOF data acquired with n=5 mice each from one representative experiment of three independent *in vivo* tumor growth experiments. **B,** Density plots for cells in each tumor type from **(A). C,** Feature plots for selected markers colored by asinh transformed staining intensity. **D,** Clustering of DA cells from **(A)** into unsupervised cell subsets. **E,** Heatmap of median marker intensities for cells in each DA cluster. nDA, non-DA cells. Colored dots indicate clusters more abundant in B16F10 (blue) or LLC (red) tumors. **F,** Manually gated CD8 T cell subset quantified for B16F10 and LLC tumors. P value by one-tailed Mann-Whitney test. **G,** DA analysis as in **(A)** for CD44+ CD4 Tconv cells. **H,** Density plots for cells in each tumor type. **I,** Feature plots for selected markers colored by asinh transformed staining intensity. **J,** Clustering of DA cells from **(G)** into unsupervised cell subsets. **K,** Heatmap of median marker intensities for cells in each DA cluster. Colored dots indicate clusters more abundant in B16F10 (blue) or LLC (red) tumors. **L,** Manually gated CD4 Tconv cell subset quantified for B16F10 and LLC tumors. P value by one-tailed Mann-Whitney test. For all applicable panels, error bars represent mean +/- SD.

Quantification of cells with high expression of Ki67, GrB, CD39, CD38, and PD1 by manual gating confirmed the relative paucity of this CD8 T cell subset in LLC MSI-H tumors (Fig. 2F).

Analysis of CD44+ CD4 Tconv cells similarly showed DA cells largely separated in UMAP space and protein expression (Fig. 2G-I). Clustering of DA cells revealed clusters 3 and 4, which were enriched in B16F10 MSI-H tumors, characterized by high expression of Ki67, PD1, CD39, and CD38 (Fig. 2J and K). Similar to the CD8 T cell subsets, cluster 2 cells enriched in LLC MSI-H tumors were TCF1+ and PD1-. Manual gating of the cell subset corresponding to CD4 clusters 3 and 4 in individual mice confirmed relative depletion in LLC MSI-H tumors (Fig. 2L). We also performed the same analysis including T cells from the ICI-responsive EO771 MSI-H and ICI-resistant EMT6 models in the ICI-treated setting. The EO771 MSI-H tumors, in comparison to the resistant LLC MSI-H or EMT6 tumors, also showed an enrichment in CD8 T cells highly expressing Ki67, GrB, CD39, CD38, and PD1 and CD4 Tconv cells highly expressing Ki67, CD39, CD38, and PD1, similar to the subsets enriched in B16F10 MSI-H tumors, when compared to EMT6 (Supplementary Fig. S4). In sum, these data show that, despite increased T cell infiltration, T cells in the ICI-resistant tumors fail to acquire the activation state and effector marker phenotypes observed in T cells in the ICI-responsive models^44^.

### IL-1 and TNFα Synergistically Drive Neutrophil Inflammation via IL1R1-Expressing Tumor Cells

Next, we focused on determining cell types and pathways responsible for the divergent T cell states in each model. In both LLC MSS and MSI-H tumors, treatment with ICIs led to higher neutrophil infiltration compared to IgG-treated controls, or B16F10 MSI-H tumors with or without treatment (Fig. 3A). Compared to mice with B16F10 MSI-H tumors, those with LLC MSI-H tumors treated with or without ICIs exhibited significantly more positive or negative pairwise frequency correlations between neutrophil cell subsets and other immune cell subsets across tumor, blood, tumor-draining lymph node and spleen (Fig. 3B), suggesting system-wide coordination we previously described for the LLC model^49^. Sorted neutrophils from ICI-treated LLC MSI-H tumors suppressed OT-I T cell proliferation *in vitro* (Fig. 3C), consistent with the function of polymorphonuclear myeloid-derived suppressor cells (PMN-MDSCs) that are extensively described in the context of cancer^50^. We subsequently examined the cytokine milieu of the TME to understand factors that can drive these changes in the LLC MSI-H tumor-bearing mice. Principal component analysis of cytokines in tumor lysates revealed that ICI treatment resulted in increased levels of cytokines regulated by the NF-κB pathway, including TNFα, IL-1β, G-CSF, and CXCL1 (Fig. 3D, Supplementary Table S3). Indeed, further quantification in tumor lysates confirmed that ICI treatment results in increased amounts of the neutrophil growth factor G-CSF, and neutrophil chemoattractants CXCL1 (Fig. 3E), and CXCL2 (Supplementary Fig. S5A). Quantitative PCR of *Csf3* (coding for G-CSF) and *Cxcl1* transcripts from sorted cells from the TME showed that tumor cells increased expression of both transcripts upon ICI treatment (Fig. 3F; Supplementary Fig. S5B and C, and Supplementary Fig. S12A). These data are consistent with our own prior data and the literature reporting production of G-CSF and CXCL1 from mouse and human tumors^49,51–54^ and suggest that ICI treatment further upregulates this existing circuit. Of note, fibroblasts in tumor were also capable of inducing G-CSF transcription following ICIs (Fig. 3F), although in the LLC TME, tumor cells outnumber fibroblasts by 10:1 (Supplementary Fig. S5D).

**Figure 3:**
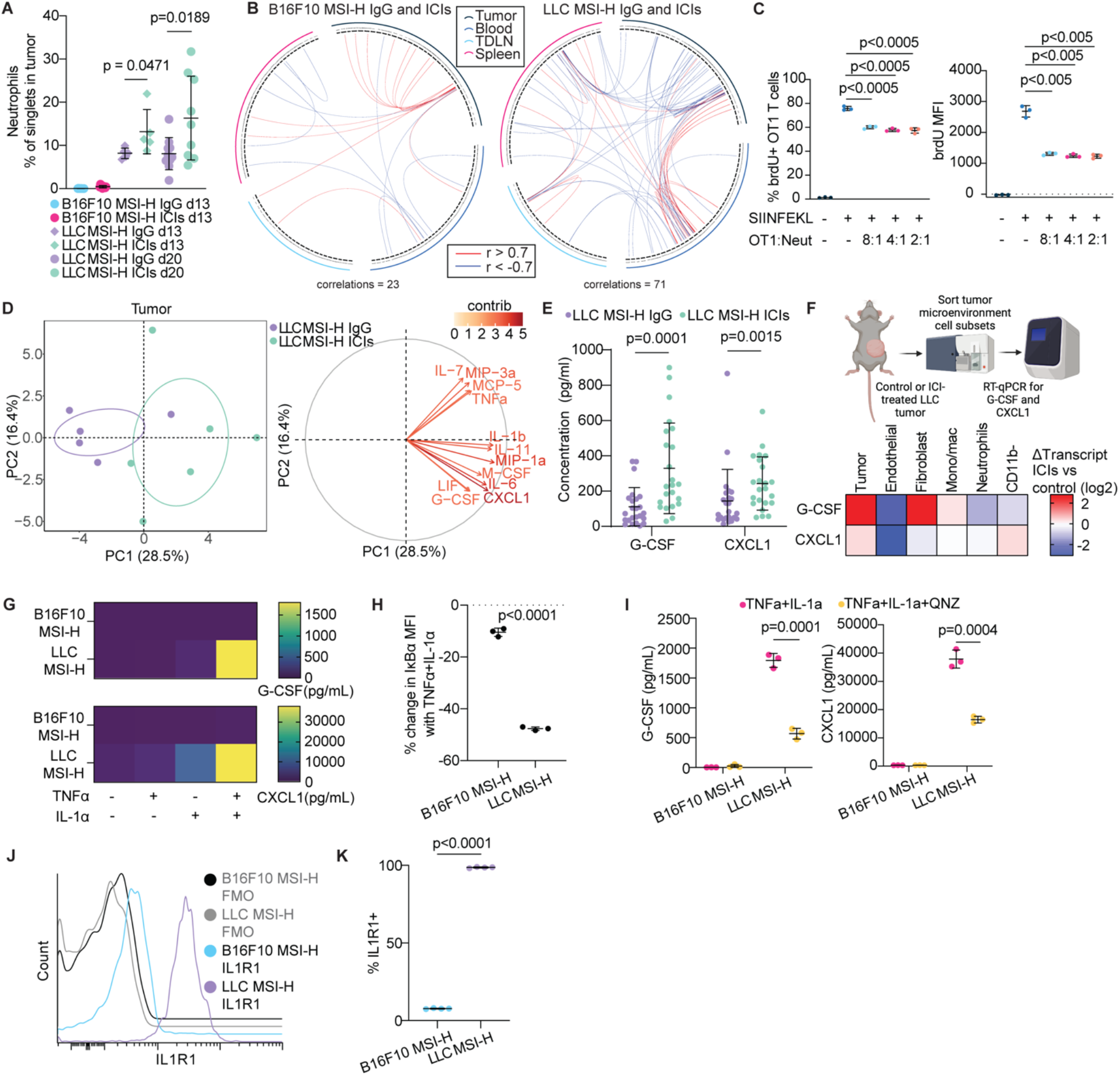
IL-1 and TNF**α** synergistically drive immunosuppressive neutrophil inflammation from IL1R1- expressing tumor cells. **A,** Quantification of neutrophils in B16F10 and LLC tumors at indicated timepoints and treatment conditions (n=5-10 mice) from two independent experiments. P values by one-tailed Welch’s t test. **B,** Pairwise Spearman correlation analysis of all unsupervised immune cell clusters across tumor, peripheral blood, tumor-draining lymph node, and spleen from CyTOF analysis of one representative experiment. Connecting lines indicate pairs of correlated clusters of which at least one cluster is a neutrophil, indicating positively (red) or negatively (blue) correlated pairs of clusters with r >0.7 or r <-0.7. Samples harvested at day 13 from n=20 mice for B16F10, and at day 20 from n=10 mice for LLC. **C,** brdU median fluorescence intensity (MFI) and % positive cell subset quantification for splenic OT-I T cells cultured with SIINFEKL peptide and/or sorted neutrophils from day 20 LLC MSI-H tumors, from n=3 mice per condition in one experiment. Adjusted p values by Welch’s t tests. **D,** Left panel, principal component (PC) analysis of cytokines in day 20 tumor lysates detected by multiplexed bead assay in one representative experiment, n=5 per condition. 95% confidence ellipse is shown. Right panel, loadings plot of contribution by the top 12 cytokines. **E,** Quantification of indicated cytokines in day 20 tumor lysates by multiplexed bead assay in 5 independent experiments. Two-tailed p values by Mann-Whitney test. **F,** RT-qPCR of cytokine transcripts from sorted cells from day 20 LLC MSI-H ICI treated tumors, with expression normalized to control treated tumors. Values represent mean of n=3-6 biological replicates from two independent experiments. **G,** ELISA quantification of G-CSF in cell culture supernatant of indicated cell lines treated with or without indicated cytokines for 24h, in one representative experiment with three biological replicates. 5 independent experiments were performed. **H,** Quantification of IκBα median fluorescence intensity of indicated cell lines with or without addition TNFα and IL-1α, in one representative experiment with three biological replicates. Two independent experiments were performed. P value by two-tailed t test. **I,** ELISA quantification of G-CSF and CXCL1 in cell culture supernatant of indicated cell lines following indicated treatment conditions from one representative experiment with three biological replicates. Two independent experiments were performed. P value by two-tailed t test. **J,** Representative histogram of IL1R1 fluorescence staining intensity for indicated cell lines in comparison to fluorescence-minus-one controls. **K,** Quantification of % IL1R1+ population in indicated cell lines from one representative experiment with four biological replicates. Two independent experiments were performed. P value by two-tailed t test. For all applicable panels, error bars represent mean +/- SD.

We previously reported that G-CSF production by LLC tumor cells is dependent on signaling through the IL-1 Receptor 1 (IL1R1)^49^. IL-1α and IL-1β, both of which can signal through IL1R1, are known to synergize with TNFα through the canonical NF-κB pathway to upregulate G-CSF, CXCL1 and CXCL2 production, possibly via IL-1-induced increase in TNF receptors^55,56^. In the TME of LLC tumors, we detected both IL-1α and IL-1β, where the former is present in larger quantities while the latter is induced upon ICIs (Supplementary Fig. S5E, F). The transcript for each cytokine increased with ICIs from a mix of tumor and immune cell types (Supplementary Fig. S5G, H). We then confirmed that LLC MSI-H cells and EMT6 cells increase G-CSF and CXCL1 production after stimulation by TNFα and IL-1α *in vitro,* in comparison to minimal responses in B16F10 and EO771 MSI-H tumor cells (Fig. 3G, Supplementary Fig. S6A-D). G-CSF and CXCL1 induction by TNFα plus IL-1α or IL- 1β were synergistic, with greater than additive responses seen with the combination (Fig. 3G, Supplementary Figs. S6A-G). Interestingly, maximal synergism for G-CSF production could be achieved with very low levels of IL-1β (1ng/mL), whereas increasing amounts TNFα potentiated the synergy (Supplementary Fig. S6E-H). Turning our attention to the downstream NF-κB pathway, LLC MSI-H cells degraded IκB more significantly compared to B16F10 MSI-H cells, consistent with greater NF-κB pathway activation in the LLC model (Fig. 3H, Supplementary Fig. S6I). Furthermore, the addition of NF-κB inhibitor QNZ suppressed G-CSF and CXCL1 expression from LLC tumor cells (Fig. 3I).

To understand why LLC and EMT6 cells were more sensitive to stimulation by IL-1 and TNFα, we examined the protein expression of IL1R1 and TNFR1 by flow cytometry. This revealed that LLC MSI-H and EMT6 cells have a high expression of IL1R1 in contrast to minimal expression in B16F10 MSI-H and EO771 MSI-H cells, while all cell lines expressed TNFR1 when compared to fluorescence-minus-one controls (Fig. 3J and K; Supplementary Fig. S7A-E). There was no difference in IL1R1 expression between LLC MSS vs. MSI-H cells (Supplementary Fig. S7F). We further investigated IL-1 and TNF pathway gene expression from bulk RNA sequencing data from LLC and B16F10 lines. For IL-1 pathway genes, we found that *Il1r1* was the most differentially regulated gene between the two models, with minimal expression of *Il1r2* and similar expression of *Il1rap* (Supplementary Fig. S7G). When comparing TNF pathway genes, we confirmed similar high expression of *Tnfrsf1a,* coding for TNFR1 (Supplementary Fig. S7H). *Tnfrsf1b* (coding for TNFR2), which can contribute to CXCL1 inflammation^57^, was expressed at an overall low level in each model but was slightly increased in LLC cells (Supplementary Fig. S7H). In sum, these data implicate an inflammatory immunosuppressive circuit supporting ICI resistance in tumor types responsive to IL-1 and TNFα stimulation, notable for their expression of IL1R1.

### IL1R1 Perturbation Limits Immunosuppressive Inflammation and Restores Favorable T Cell States to Sensitize Resistant Tumors to ICIs

We next evaluated the effect of perturbing this inflammatory axis *in vivo*. Blockade of IL-1 signaling with IL1R1-neutralizing antibody reduced G-CSF and CXCL1 in LLC MSI-H tumors (Supplementary Fig. S8A), decreased neutrophil recruitment to the tumor (Supplementary Fig. S8B), and sensitized LLC MSI-H tumors to treatment with ICIs (Fig. 4A and B; Supplementary Fig. S8C). Importantly, tumor-specific deletion of IL1R1 via CRISPR also sensitized LLC MSI-H tumors to ICIs, implicating tumor-intrinsic IL1R1 signaling during ICI resistance (Fig. 4C and D; Supplementary Fig. S8D). We additionally tested the effect of antagonizing G-CSF, downstream of IL1R1 signaling, via tumor implantation in G-CSF receptor-null mice or injection of G-CSF neutralizing antibody. This showed that loss of host G-CSF receptor signaling depletes neutrophils from tumors and improves tumor control with ICIs, while delivery of G-CSF neutralizing antibody also increases tumor control with ICIs (Supplementary Fig. S8E-J).

**Figure 4:**
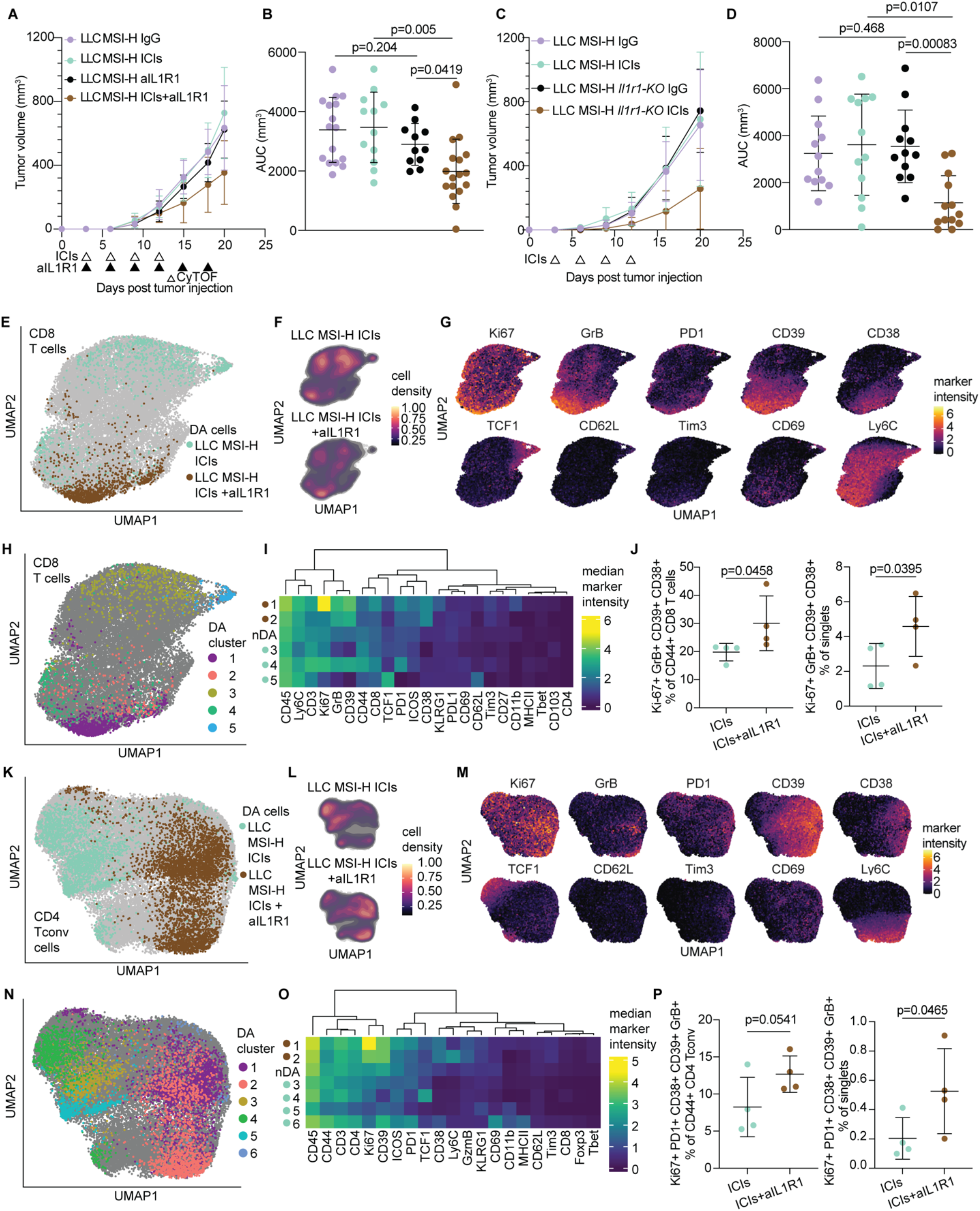
IL1R1 perturbation limits immunosuppressive inflammation to sensitize resistant tumors to ICIs. **A and B,** Quantification of mean volumes and area under curve (AUC) for LLC MSI-H tumors treated with or without ICIs and IL1R1-neutralizing antibody treatment at the indicated timepoints (arrowheads). n=11-17 mice per condition from three independent experiments. Two-tailed adjusted p value by Mann-Whitney test. **C and D,** Quantification of mean volumes and AUCs for LLC MSI-H tumors with or without tumor *Il1r1* knock out (*Il1r1*- KO) treated with or without ICIs. n=12-13 mice per condition from two independent experiments. Two-tailed adjusted p value by Mann-Whitney test. **E,** Differential abundance (DA) analysis of 26,400 CD44+ CD8 T cells from ICI-treated LLC MSI-H tumors +/- aIL1R1 treatment (n=4 mice each from one representative experiment analyzed by CyTOF) harvested at day 13. Colored dots indicate thresholded top DA cells increased in the indicated condition. **F,** Density plots for cells in each condition. **G,** Feature plots for selected markers colored by asinh transformed staining intensity. **H,** Clustering of DA cells from **(E)** into unsupervised cell subsets. **I,** Heatmap of median marker intensities for cells in each DA cluster. nDA, non-DA cells. Colored dots indicate clusters enriched in the aIL1R1 condition (brown) vs. control (teal). **J,** Manually gated CD8 T cell subset quantified for each condition, quantified as % of parental (left panel) and of singlets in tumor (right panel). P value by one-tailed t test. **K,** DA analysis as in **(E)** for CD44+ CD4 Tconv cells. **L,** Density plots for cells in each condition. **M,** Feature plots for selected markers colored by asinh transformed staining intensity. **N,** Clustering of DA cells from **(K)** into unsupervised cell subsets. **O,** Heatmap of median marker intensities for cells in each DA cluster. nDA, non-DA cells. Colored dots indicate clusters enriched in the aIL1R1 condition (brown) vs. control (teal). **P,** Manually gated CD4 Tconv cell subset quantified for each condition, quantified as % of parental (left panel) and of singlets in tumor (right panel). P value by one-tailed t test. For all applicable panels, error bars represent mean +/- SD.

Next, we performed CyTOF analyses of antigen experienced CD44+ CD8 and CD4 Tconv cells to determine changes in T cell states upon IL1R1 blockade for LLC MSI-H tumors treated with ICIs. DA analysis identified a group of cells enriched in the IL1R1 blockade condition that occupied UMAP space with higher expression of effector T cell molecules such as GrB and CD39, akin to CD8 T cells more frequently found in B16F10 tumors from the previous analysis (Fig. 4E-G; Fig. 2E). Clustering of DA cells highlighted this phenotype, identifying cluster 1, which demonstrated high median expression of Ki67, GrB, CD39, and CD38 (Fig. 4H and I). Quantification by manual gating on individual mice confirmed an increase in this cell subset in the IL1R1 blockade group (Fig. 4J). Similarly, we analyzed CD44+ CD4 Tconv cell phenotypes, identifying DA cells in the IL1R1 blockade group that expressed higher levels of effector molecules (Fig. 4K-M). Clustering the DA cells revealed clusters 1 and 2 which were enriched in the IL1R1 blockade group and demonstrating high median expression of Ki67, CD39, CD38, PD1, and GrB (Fig. 4N and O). Manual gating of this cell subset in individual mice confirmed relative enrichment with IL1R1 blockade (Fig. 4P). On the other hand, with IL1R1 blockade we found relative reductions in TCF1+ PD1- CD8 cluster 5 and CD4 cluster 4 T cells, which we previously identified as an enriched subset in LLC MSI-H tumors in comparison to B16F10 MSI-H tumors (Fig. 2E and K). Lastly, we evaluated whether IL1R1 blockade impacts other cell types within the TME in this model, such as tumor- associated macrophages or endothelial cells that can express IL1R1^58,59^. In the LLC MSI-H tumors treated with ICIs, we did not observe significant changes in cell frequencies of CD11b+ F4/80+ tumor-associated macrophages or CD31+ endothelial cells, nor significant changes in the expression of macrophage markers including PD-L1, CD206, CD38, CD39, iNOS, and MHCII (Supplementary Fig. S8K-M). These data collectively show that perturbation of the inflammatory circuit via IL1R1 inhibition sensitizes resistant tumors to ICIs and results in a shift of CD8 and CD4 T cells to cytotoxic effector-like states.

### T Cells Drive the Inflammatory ICI Resistance Circuit via TNFα

Because this inflammatory circuit was intensified by ICIs in LLC MSI-H tumors in association with T cell infiltration, and activated T cells are well-known to produce TNFα^60^, we hypothesized that T cells may paradoxically exacerbate the inflammatory resistance program when infiltrating into a TME with an active NF- κB inflammatory circuit. Depleting T cells in LLC MSI-H tumors using an anti-Thy1 antibody significantly decreased G-CSF and CXCL1 protein levels as well as neutrophil infiltration in the TME (Fig. 5A-C). Although Thy1 can be expressed in fibroblasts, they were not significantly depleted in LLC MSI-H tumors upon anti-Thy1 treatment (Supplementary Fig. S9A). Additionally, using anti-CD4 plus anti-CD8 antibodies for T cell depletion phenocopied aThy1 treatment to reduce G-CSF, CXCL1 and neutrophils, requiring elimination of both T cell types (Supplementary Fig. S9B and C). CD4 and CD8 depletion in the EMT6 model also reduced tumor- infiltrating neutrophils (Supplementary Fig. S9D). These results support a role for T cells in driving the inflammatory ICI resistance circuit.

**Figure 5:**
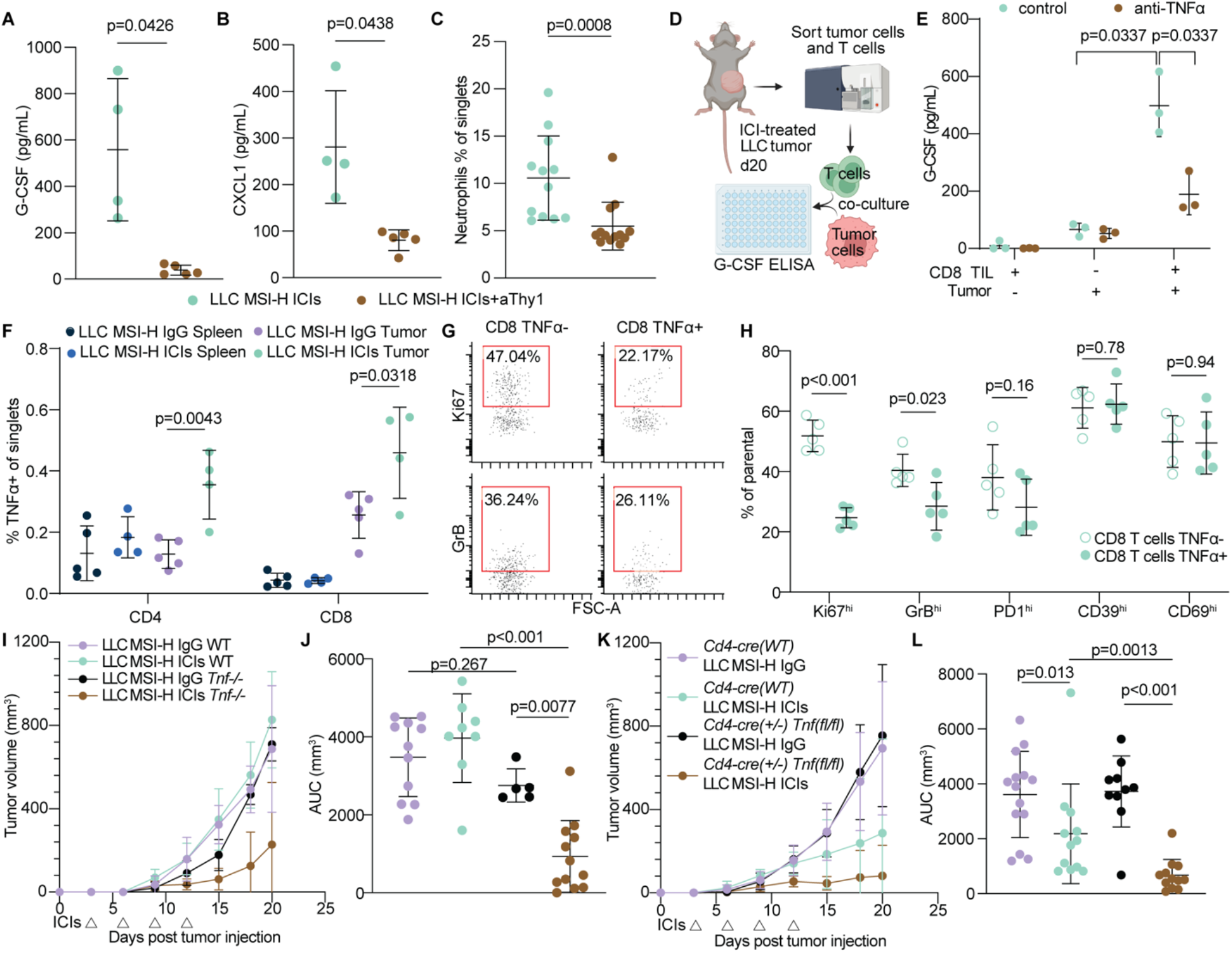
T cells drive the inflammatory ICI resistance circuit via TNFα. **A,** Quantification of G-CSF concentration in day 20 tumor lysates for indicated treatment conditions with n=4-5 mice from one experiment. P value by two-tailed Welch’s t test. **B,** Quantification as in (**A)** for CXCL1. **C,** Quantification of neutrophils in day 20 tumors for indicated treatment conditions in two independent experiments, n=12-13 mice. P value by two-tailed Welch’s t test. **D,** Experimental schematic for T cell-tumor coculture. **E,** Quantification of G-CSF concentration in supernatant of 24h culture containing indicated sorted cell types from day 20 LLC MSI-H tumors, with or without TNFα neutralizing antibody, from one representative experiment with n=3 mice. Two independent experiments were performed. Adjusted p values by two-tailed Welch’s t test. **F,** Quantification of TNFα+ CD4 or CD8 T cells in tumor or spleen of day 20 LLC MSI-H tumor bearing mice treated with indicated antibodies, n=4-5 mice from one representative experiment. Three independent experiments were performed. P value by two-tailed t test. **G,** Biaxial plots of Ki-67 and GrB fluorescence staining intensity in TNFα- or TNFα+ CD8 T cells. **H,** Quantification of TNFα- and TNFα+ CD8 T cells as % of cells binned into high expression gates for each marker. P value by two-tailed t tests. n=5 mice from one representative experiment. Two independent experiments were performed. **I and J,** Quantification of mean volumes and area under curves (AUC) for LLC MSI-H tumors treated with IgG or ICIs, grown in wild-type C57BL/6J mice or *Tnf*-null mice. Adjusted p values by Mann-Whitney test. n=5-12 from two independent experiments. **K and L,** Quantification of mean volumes and area under curves (AUC) for LLC MSI-H tumors treated with IgG or ICIs, implanted in *Cd4-cre(WT) Tnf(fl/fl or fl/+)* control mice or *Cd4-cre(+/-) Tnf(fl/fl)* mice. Adjusted p values by Mann-Whitney test. n=10-14 mice from two independent experiments. For all applicable panels, error bars represent mean +/- SD.

We next sought to directly test the interaction between T cells and tumor cells, hypothesizing that T cell- derived TNFα can stimulate tumor cells to produce inflammatory cytokines that support neutrophils. Co-culturing CD3/CD28-stimulated CD8 T cells sorted from LLC MSI-H tumors together with LLC MSI-H tumor cells increased G-CSF production as quantified in the supernatant by more than five-fold, indicating that CD8 T cells are sufficient to drive this circuit (Fig. 5D and E; Supplementary Fig. S12B). Importantly, the increase in G-CSF was largely dependent on TNFα, as the addition of a neutralizing antibody to TNFα resulted in a significant reduction of G-CSF (Fig. 5E). Neutralization of other potential T cell factors, such as FasL or GM-CSF, did not result in reduction of G-CSF (Supplementary Fig. S9E). *In vivo*, treatment with ICIs specifically increased TNFα- expressing CD4 and CD8 T cells in LLC MSI-H tumors compared to the spleen (Fig. 5F). This was also accompanied by ICI-induced increases in TNFα+ tumor-infiltrating neutrophils, a cell type well-known to express TNFα^61^ (Supplementary Fig. S9F). CD4 and CD8 depletion decreased TNFα+ neutrophils in the TME (Supplementary Fig. S9G), demonstrating dependence on T cell activity. Further characterization of TNFα+ CD8 T cells revealed that these cells are less likely to express Ki67 or GrB in comparison to TNFα- cells (Fig. 5G and H), segregating these key effector functions into different T cell subsets. Analysis of TNFα+ CD4 T cells also showed lower expression of Ki67, though these cells demonstrated higher expression of CD39 (Supplementary Fig. S9H).

To test the functional contribution of TNFα to the inflammatory ICI resistance circuit, we implanted LLC MSI-H tumors in *Tnf* knockout mice lacking host-derived TNFα, which sensitized the tumors to ICI treatment accompanied by a decrease in neutrophils (Fig. 5I and J; Supplementary Fig. S9I and J). Furthermore, we generated T cell-specific *Tnf* knockout mice as previously described, using *Cd4*-cre to delete the *Tnf(fl/fl)* allele^61^ (Supplementary Fig. S9K-M). Importantly, tumor growth was also restricted following ICIs in mice specifically lacking T cell-derived TNFα (Fig. 5K and L; Supplementary Fig. S9N). We further evaluated the impact of IL1R1 blockade in the setting of T cell TNFα loss, hypothesizing that, given the synergy between IL-1 and TNFα, we would not observe an additive benefit with the combined perturbations. Indeed, tumor control with ICIs with the combination of IL1R1 blockade and T cell TNFα knockout was not significantly different from the benefit with single perturbation alone (Supplementary Fig. S9O-Q). Collectively, these data support a novel mechanism whereby ICI-induced infiltration of T cells producing TNFα exacerbates NF-κB-driven transcription of G-CSF and CXCL1 in tumor cells to create a paradoxical circuit of immunosuppression.

### T cell, Tumor, and Neutrophil Inflammation Circuit Is Active in Human Cancer and Is Associated with Poor Clinical Outcome

We next interrogated our model of T cell-exacerbated tumor inflammation in human cancer. First, we tested whether G-CSF and CXCL1 production can be elicited from human cancer cells. We incubated breast cancer organoids derived from two different resected primary tumors (TORG40, isolated from triple-negative inflammatory breast cancer, and TORG139, isolated from triple-negative breast cancer) with IL-1α and TNFα. The TORG40 organoids demonstrated significant induction of G-CSF and CXCL1 secretion in the supernatant, while the TORG139 organoids did not produce G-CSF and had a diminished induction of CXCL1 in comparison (Fig. 6A and B). RT-qPCR showed that TORG40 organoids also had a higher induction of *CXCL8* transcripts compared to TORG139 (Supplementary Fig. S10A). Based on our mouse data, we hypothesized that TORG40 cells expresses *IL1R1* more highly compared to TORG139 cells. Indeed, TORG40 cells had higher *IL1R1* but similar *TNFR1* expression compared to TORG139 cells (Fig. 6C; Supplementary Fig. S10B). These data further support our findings in mouse tumor cells that IL1R1 expression distinguishes tumors sensitive to IL-1 and TNFα stimulation.

**Figure 6:**
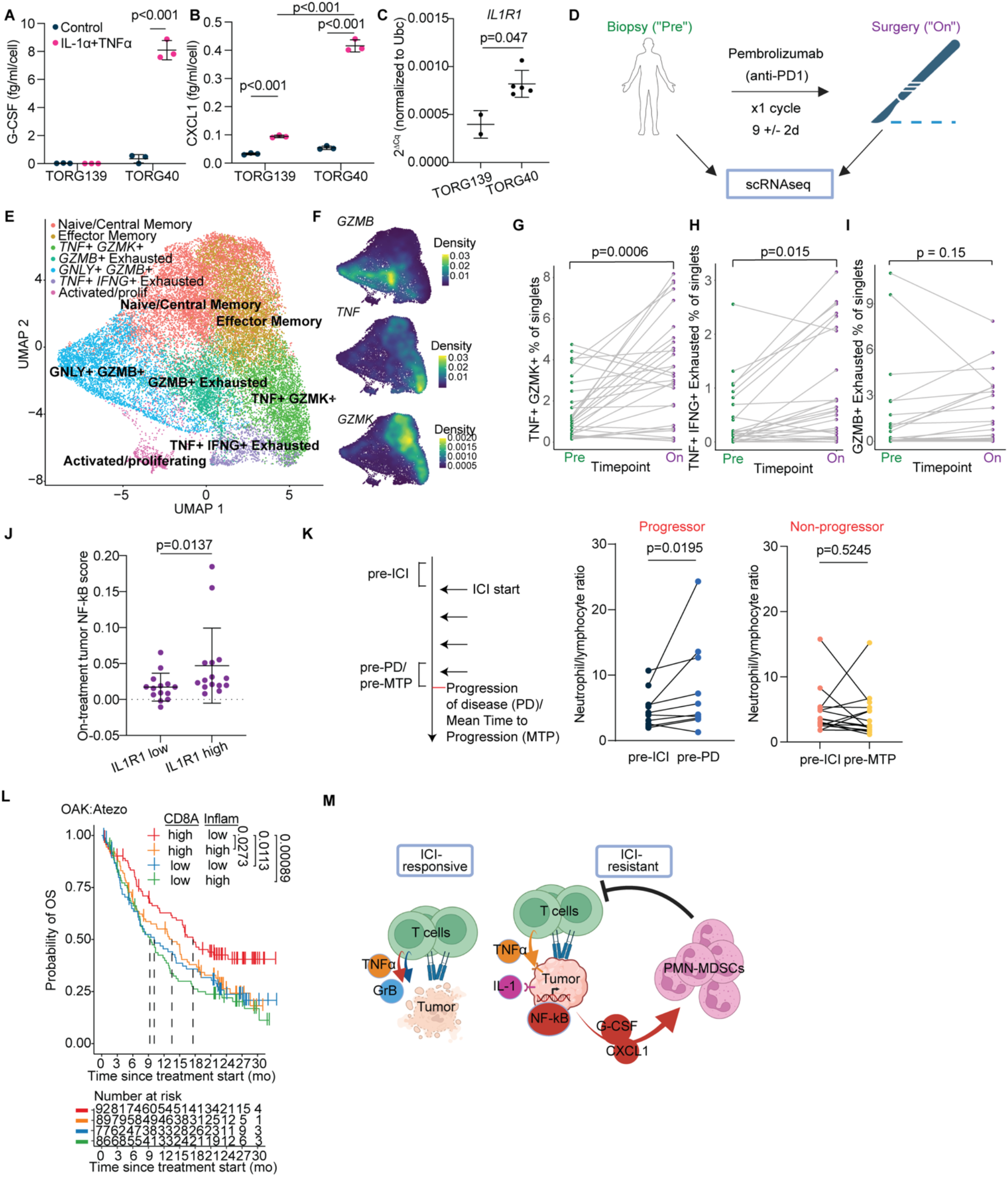
T cell, tumor, and neutrophil inflammation circuit in human cancer. **A,** ELISA quantification of G-CSF in supernatant of breast cancer organoid culture following indicated treatments for 24h, from one representative experiment with n=3 biological replicates. Two independent experiments were performed. P value by two-tailed t test. **B,** Analysis as in **(A)** for CXCL1. Adjusted p values by t tests. **C,** *IL1R1* transcript quantification by RT-qPCR for breast organoid models from two independent experiments. P value by one-tailed Mann-Whitney test. **D,** Schematic of the BIOKEY study (EGAS00001004809) analyzed in **(E-J). E,** UMAP of 24,349 CD8 T cells color coded by CD8 T cell phenotype. **F,** UMAP cell density plots of *GZMB*, *TNF*, and *GZMK* expressing cells. **G-I,** Percent of cells in indicated cluster out of singlet cells in tumor for pre vs. on- treatment samples for each patient. P value by two-tailed Wilcoxon matched-pairs signed rank test. **J,** Quantification of on-treatment NF-κB scores in tumor cells for each patient, stratified by *IL1R1* expression in ≥1% (high) or <1% (low) of tumor cells. **K,** Definitional schema and plot of neutrophil/lymphocyte ratios in peripheral blood of patients with MSI-H tumors treated with ICI therapy, stratified by progression vs. non-progression status. P value by two-tailed Wilcoxon matched-pairs signed rank test. **L,** Kaplan-Meier curves of patients in the OAK trial (NCT02008227), stratified by *CD8A* status and inflammation gene score status (“inflam”). P values by log- rank test. **M,** Mechanistic model.

To understand the relationship between T cells and tumor cells in more detail in the context of ICI treatment, we analyzed a publicly available single-cell RNA sequencing dataset of tumor samples from non- metastatic breast cancer patients before and after a single dose of pembrolizumab^62^ (Fig. 6D). A total of 7 clusters were identified across all CD8 T cells (Fig. 6E; Supplementary Fig. S10C). Notably, we observed a separation of CD8 T cells expressing *TNF* from those expressing GrB (*GZMB),* or Ki67, which was expressed by cells in the Activated/proliferating cluster (Fig. 6F; Supplementary Fig. S10C), paralleling the results from our mouse models (Fig. 5G and H). Comparison of differentially expressed genes in the two *TNF-*expressing clusters *– TNF+GZMK+* and *TNF+ IFNG+* exhausted – to the *GZMB+* exhausted cluster revealed higher expression of markers suggestive of activation (*CD69*^63^*, FOS*^64^*, DUSP1*^65^, and *CCL4*^66^; Supplementary Fig. S10D and E). Importantly, we found that the *TNF*+ *GZMK*+ cluster and *TNF+ IFNG+* exhausted cluster significantly increased in patients following pembrolizumab treatment, whereas the *GZMB*+ Exhausted cluster was not significantly changed (Fig. 6G-I, Supplementary Fig. S10F). The increases in the *TNF*-expressing clusters were positively correlated with each other (Supplementary Fig. S10G). We additionally examined *TNF* expression from CD4 T cells and myeloid cells in this dataset. For CD4 T cells, a variety of subsets expressed *TNF,* ranging from Th0, Th1, and Th17 cells (Supplementary Fig. S11A-C). Among these, *TNF*+ Th1 subsets increased following ICI treatment, characterized by expression of *ENTPD1* (coding for CD39), which we previously noted to be expressed highly by TNFα+ mouse CD4 T cells (Supplementary Figs. S11D and E, and S9H). For myeloid cells, a portion of C1q+ macrophages expressing *TNF* increased following treatment (Supplementary Fig. S11F-I).

Next, we hypothesized that in the setting of an increase in *TNF-*expressing cells with ICIs, *IL1R1-*expressing tumors would preferentially activate tumor-intrinsic NF-κB inflammation. For tumor cells in each patient sample, we determined the level of *IL1R1* expression as well as tumor cell-specific score of NF-κB activity based on an established gene set^67^. Tumor NF-κB scores were higher in the IL1R1-high group of patients following ICIs (Fig. 6J). Furthermore, increases in the CD8 *TNF+ GZMK+* and *TNF+ IFNG+* exhausted cluster frequencies were positively correlated with increases in tumor NF-kB scores in the IL1R1-high group, but not in the IL1R1-low group (Supplementary Fig. S11J). These results demonstrate that ICI treatment can induce an expansion of inflammatory leukocytes characterized by *TNF* expression that are associated with increased tumor- intrinsic NF-κB signaling when the tumor cells express *IL1R1*.

Because droplet-based single-cell RNA-sequencing does not efficiently capture neutrophils^68^, we sought to test our hypotheses regarding neutrophils using additional human datasets containing neutrophil quantification. Recently, a multicellular and molecular inflammatory hub composed of NF-κB transcription factors, inflammatory tumor cells, lymphocytes, neutrophils, and neutrophil-attracting chemokines has been described in human colorectal cancer (COAD)^69^. We hypothesized that within a TME capable of upregulating such NF-κB- dependent, neutrophil-recruiting inflammation, MSI-H status and the accompanying increased T cell infiltration^70,71^ would result in an increase in neutrophils in the TME. Inference of TME immune composition using CIBERTSORTx^72^ for analysis of the Cancer Genome Atlas (TCGA) RNA sequencing dataset for COAD patients, grouped by their MSI status, showed an increase in neutrophils in a subset of MSI-H tumors compared to MSS tumors (Supplementary Fig. S11K). We reasoned furthermore that, given the systemic mobilization of neutrophils during ICI resistance in antigen-rich tumors in mice, ICI resistance in patients with MSI-H cancers may be associated with exacerbated systemic neutrophilia after treatment. We identified MSI-H cancers in our institutional clinical next-generation sequencing database and analyzed those who were started on ICI therapy but experienced treatment failure. When compared to a 6-week period prior to initiation of ICIs, the peripheral blood neutrophil to lymphocyte ratio (NLR) was increased after ICI treatment in the period preceding disease progression (Fig. 6K). On the other hand, for patients who did not experience progression, NLR did not change over the same interval of time. NLR has been associated with poor prognosis in various human cancer types, including but not exclusive to MSI-H cancers^73,74^.

Lastly, we determined the significance of the T cell-tumor-neutrophil inflammatory circuit with regard to clinical outcomes for ICI-treated patients. To this end, we analyzed RNA sequencing data from the atezolizumab arm of the OAK trial (NCT02008227), which was a randomized phase III trial comparing atezolizumab vs. docetaxel in locally advanced or metastatic non-small cell lung cancer patients^75,76^. Following stratification of patients into *CD8A* high vs. low groups, we determined the impact of a custom gene set score “inflam” consisting of genes involved in the ICI resistance circuit we have described, including *IL1R1, TNFRSF1A, IL1A, IL1B, TNF, CD69, CSF3, CXCL1, CXCL2, CXCL8, NFKB1, NFKB2.* This analysis demonstrated that only patients in the CD8A high but “inflam” low group had superior overall survival compared to the other groups in aggregate (p=0.00124; HR=1.667 (1.222-2.278)), as well as to each individual group (CD8A high, “inflam” high: p=0.0273, HR=1.517 (1.048-2.196)); CD8A low, “inflam” low: p=0.0113, HR=1.634 (1.117-2.392); and CD8A low, “inflam” low p=0.00089, HR=1.8642 (1.291-2.692)) (Fig.6L). Notably, those patients with CD8A high and “imflam” high tumors, similar to our resistant mouse models, did not experience a survival benefit. Analysis of progression-free survival (PFS) demonstrated similar trends, with patients in the CD8A high but “inflam” low group exhibiting the most favorable outcomes (Supplementary Fig. S11L). These data provide additional evidence in support of the proposed ICI resistance mechanism in tumors with T cell infiltration (Fig. 6M).

## Discussion

In this study, we define a mechanism by which T cells exacerbate an inflammation circuit involving NF-κB pathway activation in IL1R1+ tumor cells, paradoxically driving an immunosuppressive TME characterized by neutrophil recruitment and resistance to ICI therapy. NF-κB is a signaling pathway activated in many cell types in the TME and is known to both promote and prevent tumor progression depending on context^19^. NF-κB signaling in cancer and myeloid immune cells, however, frequently results in pro-tumor inflammation during tumorigenesis and progression^77,78^. Notably, inhibition of TNFα has recently been shown to improve ICI efficacy in mouse models, in addition to protecting against autoimmune adverse effects^35,36^. However, TNFα can be produced by many cell types^61^, and therefore it has remained unclear to what extent T cells contribute to the effects of TNFα inhibition against cancer. Our data show that T cell-derived TNFα can potentiate the production of NF-κB-regulated soluble factors that promote adaptive resistance to therapy in IL1R1-expressing tumors.

Prior publications have highlighted the importance of an immunosuppressive axis involving immunosuppressive neutrophils (PMN-MDSCs) that are critically relevant for cancer treatment outcome^50^. Notably, the Melero and Datta groups have recently reported IL-1 and TNF can promote CXCL1, CXCL2 and IL-8 (in humans) from tumor cells and fibroblasts, culminating in PMN-MDSC recruitment to the TME^57,79^. Our results build on these recent findings, and we now provide new functional data demonstrating a paradoxical role for T cell-derived TNFα in exacerbating this inflammatory resistance circuit after ICI treatment. In cancer immunity, various T cell subsets have been implicated in tumor development and progression, most notably Tregs, which classically suppress effector T cell responses^80^. Th17 CD4, as well as γδ T cells, can also drive pro-tumor inflammation via IL-17 production^81,82^, while CD8 T cells have been previously shown to promote chemical carcinogenesis^83,84^. Interestingly, in the context of autoimmunity and immune-related adverse events, effector memory CD4 T cells have been recently shown to instruct NF-κB inflammation in partnering dendritic cells in a TNFR- and CD40-dependent manner^85,86^. We now show that ICI treatment can result in CD4 and CD8 T cell- exacerbated immunosuppressive inflammation, mediated by T cell-derived TNFα, which synergizes with IL-1 present in the tumor microenvironment to limit therapeutic efficacy. This adds nuance to the prevailing view that T cell effector functions are beneficial in the TME and for response to immunotherapy, distinguishing productive from detrimental effector functions that depend on the context of the tumor. Our data importantly demonstrate that tumor cells expressing IL1R1 can best participate in the immunosuppressive circuit due to synergistic activation of NF-κB by IL-1 and TNFα.

While we have focused on the role of IL-1 and TNFα in signaling through tumor cells to elicit immunosuppressive neutrophilic inflammation, it is important to note that there may be a number of complementary mechanisms through which these cytokines amplify immune suppression in the TME. IL1R1 expression on host cells, including fibroblasts, has been shown to contribute to accumulation and potentiation of MDSC activity^87,88^. In addition, TNFα can act directly on cells in the TME to promote tumor progression, including induction of T cell activation-induced cell death^89,90^, tumor cell dedifferentiation^91^, and stabilization of immunosuppressive molecules such as PD-L1, CD73, CD39, and TGF-β, which can potentiate CD4 Treg function^33,92^. Our own data specifically provide support for the function of tumor cell-expressed IL1R1 and T cell-derived TNFα toward ICI efficacy, and additional experiments are needed to dissect the contribution and interaction of each of these intersectional mechanisms.

While we have leveraged T cell-infiltrated MSI-H tumors for our study, the mechanism presented is not specific to MSI-H tumors. We show the ICI resistance circuit is also active in the EMT6 breast cancer model (which does not exhibit MSI), breast cancer organoids, and in non-MSI breast and lung cancer. Rather than MSI status, the common denominators for engaging the resistance circuit are recruitment of TNF-expressing T cells, presence of IL-1, and expression of IL1R1 on tumor cells, which collectively result in activation of the NF-κB pathway to drive neutrophil recruitment. In patients, our results indicate that neutrophil predominance as quantified by NLR is associated with ICI resistance in MSI-H tumors and that a gene signature of this resistance circuit is correlated with ICI treatment outcome in lung cancer patients, whose tumors rarely display MSI-H status^93^. We hypothesize that the increased antigenic burden in MSI-H tumors can exacerbate T cell-mediated inflammation in tumors responsive to IL-1 and TNFα, although we do not rule out potential contribution from immune-stimulatory effects of genome instability induced by MMRd^94,95^.

Importantly, our study revises the prevailing paradigm of ICI efficacy and offers a potential explanation for why patients may experience poor responses to ICIs despite predicted or confirmed T cell infiltration, beyond T cell exhaustion. Importantly, while high T cell infiltration or proxy measures for T cell inflammation such as high tumor mutational burden or MMRd/MSI-H status generally correlate with a relative improvement in ICI responses as compared to non-T cell inflamed controls, it is notable that often, the majority of patients in the high T cell inflammation group still do not respond to therapy^11,12,96^. Our findings now refine the current models of ICI response to include the active role that tumor-infiltrating CD4 and CD8 T cells can play in shaping the inflammatory tone of the TME, including unexpectedly exacerbating an immunosuppressive tissue environment.

Strategies targeting the IL-1/IL1R1/TNF axis are being tested in clinical trials. Based on the finding that IL-1β inhibition via Canakinumab may have reduced lung cancer incidence and mortality, a series of phase III CANOPY clinical trials were launched and recently reported their results, albeit demonstrating no significant impact on survival outcomes during treatment of non-small cell lung cancer with inhibition of IL-1β alone^97–100^. A phase I-II trial is ongoing to test the safety and efficacy of an IL1R1 inhibitor Isuanakinra in combination with ICIs in patients with advanced solid tumors (NCT04121442). The combination of TNF inhibitors Infliximab or Certolizumab with Nivolumab and Ipilimumab was recently tested in the TICIMEL phase Ib trial in patients with advanced or metastatic melanoma, with a good safety profile and promising response rate^101^. Lastly, an inhibitor of CXCR1 and CXCR2, which are receptors for neutrophil chemokines including CXCL1, CXCL2, and CXCL8, is being tested in combination with ICIs in patients with melanoma, non-small cell lung cancer, and colorectal cancer (NCT03161431, NCT05570825, NCT04599140). Our results contribute to the rationale for these ICI combination approaches, and further suggest that a subset of patients who could particularly benefit from ICIs and IL1R1/TNFα blockade may be those with T cell-infiltrated tumors and IL1R1^+^ tumor cells. Lastly, we provide evidence for G-CSF/G-CSFR inhibition as a potential therapeutic strategy in combination with ICIs, an approach that has not yet entered clinical trials.

In summary, our data elucidate a mechanism by which T cells can unexpectedly promote ICI resistance in a manner dependent on the tumor context. These results can inform new strategies for therapy selection and immune monitoring as well as therapeutic strategies to convert inflammation-promoting T cell activity into productive anti-tumor immunity.

## Methods

### Cell lines

Parental LLC cells (RRID:CVCL_4358) were gifted by Dr. Ross Levine (Memorial Sloan Kettering Cancer Center). B16F10 cells (RRID:CVCL_0159) were gifted by Dr. Jeffrey Bluestone (University of California, San Francisco (UCSF)). EO771 cells (RRID:CVCL_GR23) were gifted from Dr. Robin Anderson (Peter MacCallum Cancer Center). EMT6 cells (RRID:CVCL_1923) were purchased from ATCC. LLC, B16F10, and EO771 parental and derivative cell lines were cultured in DMEM supplemented with 10% fetal calf serum, 10mM HEPES, and 100 U mL^-1^ penicillin and 100 μg mL^-1^ streptomycin. EMT6 cell lines were cultured in Waymouth’s MB 752/1 medium with 2mM L-glutamine, 15% fetal calf serum and 100 U mL^-1^ penicillin and 100 μg mL^-1^ streptomycin. For generation of MSH2 and IL1R1 knock out cell lines, a combination of two targeting guide crRNAs (Dharmacon), or scrambled nontargeting control, were mixed with tracrRNA and recombinant Cas9 protein (QB3 Macrolab, UC Berkeley) at 1:1:2:2 molar ratio and incubated at 37°C to form the ribonucleoprotein complex. This was then mixed with 1ul of 100uM single-stranded oligodeoxynucleotides (ssODN) enhancer, and 3 x 10^5^ trypsinized cells in electroporation buffer and electroporated using the Lonza 4D 96-well electroporation system. Nucleic acid sequences, electroporation buffer type and electroporation code are found in Supplementary Table 1. After a 15 minute recovery in a 37°C CO2 incubator, cells were expanded in culture at subculture ratios of 1:10-1:15 every 48 hours. MSI-H cell lines used have undergone 60 passages in total. All cell lines underwent mycoplasma testing with negative results as of November 1, 2023. Cells were used for experiments within 2-3 passages of thaw.

Following the initial experiments included in this manuscript, cell line verification testing was performed via short tandem repeat (STR) analysis on the syngeneic mouse tumor cell lines used. All cell lines were successfully matched, but it revealed that a cell line we had previously attributed to be the AT3 breast cancer cell line (RRID:CVCL_VR89) was in fact the LLC cell line (RRID: CVCL_4358), along with its MSS and MSI-H derivative lines^49^. To further ensure that the cell line was correctly identified as LLC, we performed a PCR assay for PyMT transgene detection, which should be present in the AT3 line because it was derived from an MMTV- PyMT mouse^102^. We confirmed this to be true in an independently purchased reference AT3 cell line, now available from Millipore, but not in our LLC cell lines (Supplementary Figs. S2, and S13). Furthermore, we used exome sequencing data from our cell line to verify the presence the of two point mutations reported in the literature to be found in the LLC line^103,104^ (Supplementary Table S4).

### Animals and *in vivo* treatments

All mice were housed in an American Association for the Accreditation of Laboratory Animal Care- accredited animal facility and maintained in specific pathogen-free conditions. Animal experiments were approved by and conducted in accordance with Institutional Animal Care & Use Program protocol AN184195. Female C57BL/6J (RRID:IMSR_JAX:000664) and BALB/cJ (RRID:IMSR_JAX:000651) mice between 8 and 12 weeks old were purchased from the Jackson Laboratory. B16F10 (1 x 10^5^ cells per 100 μL), LLC (5 x 10^5^ cells per 100 μL), EO771 (8 x 10^6^ cells per 100 μL), and EMT6 (5 x 10^5^ cells per 100 μL) cells in serum-free DMEM were transplanted into the right subcutaneous flank for B16F10 melanoma, the left 4^th^ mammary fat pad for EO771 and EMT6, and either the flank or the mammary fat pad for LLC. Unless indicated as in Figure 1 and Supplementary Fig. S1 for subcutaneous implantation, LLC cells were implanted into the mammary fat pad for all experiments. Different cell numbers for inoculation were chosen for each model to address disparities in *in vivo* tumor growth rates. TCR transgenic OT-I CD45.1 mice (RRID:IMSR_JAX:003831) were bred at our facility. Female B6.129S-*Tnf^tm1Gkl^*/J mice (RRID:IMSR_JAX:005540, *Tnf-/-* in manuscript), and B6.129X1(Cg)- *Csf3r^tm1Link^*/J mice (RRID:IMSR_JAX:017838, *Csf3r-/-* in manuscript) between 7-8 weeks of age were purchased from the Jackson Laboratory. For T cell-specific *Tnf* knockout mice, *Tnf(fl/fl)* mice^61^ crossed to the Ai14 Cre reporter (RRID:IMSR_JAX:007914) were obtained (gift from Dr. Ophir Klein); these mice were crossed to *Cd4*- cre mice (RRID:IMSR_JAX:022071) to obtain T cell-specific *Tnf* knockout mice. Control *Cd4-cre(WT)* mice included mice with *Tnf(fl/fl)* or *Tnf(fl/+)* genotypes. All mice were heterozygotes or homozygotes for the Ai14 Cre reporter allele. Animals were housed under standard specific pathogen-free conditions with typical light/dark cycles and standard chow.

For all anti-PD-1 and anti-CTLA4 antibody treatments, 250 μg of anti-PD-1 clone RMP1-14 (Bio X Cell, RRID:AB_10949053) and 200 μg of anti-CTLA4 clone 9H10 (Bio X Cell, RRID: AB_10950184) in 100 μL PBS were injected intraperitoneally at days 3, 6, 9 and 12 post tumor injection. For IL1R1 inhibition, 200 μg of antibody clone JAMA-147 (Bio X Cell, RRID: AB_2661843) was injected intraperitoneally every 3 days starting at day 3 post tumor injection. For Thy1 depletion, 200 μg of antibody clone M5/49.4.1 (Bio X Cell, RRID: AB_1107681) was injected intraperitoneally at days −2, 0, 7 and 14 relative to tumor injection. For anti-CD4 (RRID: AB_1107636) and anti-CD8 (RRID: AB_1125541) antibody treatments, 400 μg of each antibody was injected intraperitoneally on days −1, 3, 7, 10, 13, and 16 post tumor injection. Where indicated by “IgG”, isotype- matched control IgG antibodies were used at the same concentrations as experimental antibodies (RRID: AB_1107769, AB_1107782). For G-CSF neutralization, 10 ug of antibody (R&D Systems MAB414, RRID: RRID:AB_2085954) was injected intraperitoneally starting at day 5 post tumor implantation.

### MSI PCR quantification

Microsatellite instability in the mouse cell lines was quantified using a panel detecting four microsatellite markers^43^. Genomic DNA from cell pellet was isolated using the GeneJET Genomic DNA Purification Kit (Thermo Fisher Scientific, K0722) according to manufacturer protocol. 20 ng of genomic DNA was used as input for a PCR reaction using Platinum Taq (Thermo Fisher Scientific, 10-966-018) with 2mM MgCl2, 0.2 μM each primer (Supplementary Table 2), and 200 μM dNTP. The cycling profile included: 1 cycle at 94 °C for 4 minutes, 94 °C for 30 seconds, 56 °C for 45 seconds and 72 °C for 30 seconds for a total of 35 cycles, and a final extension at 72 °C for 6 minutes. Samples were submitted for fragment analysis by Genewiz, and analyzed using PeakScanner 2 (Thermo Fisher Scientific).

### Next generation sequencing and neoantigen priority score analysis

Whole exome and bulk RNA sequencing were performed with flash frozen cell pellet submitted to MedGenome (Foster City, CA). For whole exome sequencing, the library was prepared using the Agilent SureSelectXT Mouse All Exon Kit, and sequenced on NovaSeq 6000 with paired-end 100 read length. Data was processed using Fastq preprocessing using FastQC v0.11.8 and adapter trimming with fastq-mcf v1.05. 99.66% of the total reads aligned to the reference genome. 87.77% of the aligned reads passed the alignment filter (mapping quality ≥ 29 and insert size between 100-1000). Reads after duplicate removal, indel realignment and recalibration featured more than 98.49% coverage and 86.67-114.47X average depth. Somatic variant calling was performed using Strelka^105^ (v2.9.10). Default settings were used for variant calling. For realignment and base- recalibration, dbsnp150 variants were used. The identified somatic variants were further filtered and only passed and on-target variants were considered for downstream analysis. Default parameters provided by Strelka were used to filter passed somatic variants. The on-target variants were filtered based on the coordinates of the target regions provided by the vendor. RNA sequencing was performed using two biological replicates per cell line using Illumina TruSeq Stranded Total RNA library preparation, and sequenced using NovaSeq with target of 50 million total reads per sample. 86.18-91.46% of total reads for each sample had Quality scores of > 30. Kallisto^106^ (v0.46.1) was used to quantify abundances of the transcripts in transcripts per million (TPM) with the strand specific read processing argument ‘--rf-stranded’.

For neoantigen priority score analysis, the MuPeXI pipeline was used as described^44^. The somatic allele frequency estimate was extracted from the strelka vcf records following the Strelka user guide and added to the vcf files. The vcf files with allele frequency, the mean expression output files from Kallisto and the GRCm38 references were given as input to the MuPeXI pipeline (v1.2.0) with the species option set to mouse and the HLA alleles set to ‘H-2-Kb,H-2-Db’.

### Mass cytometry antibodies

All mass cytometry antibodies and staining concentrations are listed in Supplementary Table S5. Primary conjugates of mass cytometry antibodies were prepared using the Maxpar antibody conjugation kit (MCP9 for Cadmium or X8 for non-Cadmium metals; Fluidigm) according to the manufacturer’s recommended protocol. After labeling, antibodies were diluted in HRP-Protector Peroxidase Stabilizer (Candor) for Cadmium conjugated antibodies, and Candor PBS Antibody Stabilizer supplemented with 0.02% sodium azide for non-Cadmium conjugated antibodies. Antibodies were stored at 4°C long term. Each antibody clone and lot were titrated to optimal staining concentrations using primary mouse samples.

### Cell preparation

Tissue preparations from tumor, tumor-draining lymph node, spleen and lymph node were performed simultaneously. After euthanasia by CO2 inhalation, peripheral blood was collected via cardiac puncture and transferred into heparin-coated vacuum tubes. Blood was diluted into ACK lysis buffer, incubated for 3 minutes on ice then quenched with 10 mL PBS with 5 mM ethylenediaminetetraacetic acid (EDTA) and 0.5% bovine serum albumin (BSA) (PBS/EDTA/BSA). Spleens and tumor-draining inguinal lymph nodes were homogenized with a syringe pusher over 70 μm filter with PBS/EDTA. Tumors were finely minced and digested in RPMI-1640 with 4 mg ml^-1^ collagenase IV (Worthington) and 0.1 mg ml^-1^ DNaseI (Sigma) in a scintillation vial with a micro stir bar for 30 minutes at 37 °C. After digestion, cells were filtered twice over 70 μm filters with PBS/EDTA. All tissue samples were centrifuged at 500 *g* for 5 minutes at 4°C, resuspended in PBS/EDTA, and mixed 1:1 with PBS/EDTA containing 100 mM cisplatin (Enzo Life Sciences) for 60 seconds before quenching 1:1 with PBS/EDTA/BSA. Cells were centrifuged again, resuspended in PBS/EDTA/BSA at a density between 1 x 10^6^ and 1 x 10^7^ cells per mL, fixed for 10 minutes at room temperature using 1.6% formaldehyde (final), washed twice with PBS/EDTA/BSA and frozen in 100ul buffer at −80°C. All samples were processed on ice with buffers at 4 °C.

### Mass cytometry

2 x 10^6^ cells from tumor and 1 x 10^6^ cells from other organs from each animal were barcoded with distinct combinations of stable Pd isotopes in 0.02% saponin in PBS as previously described^107^. Cells were washed twice with cell-staining media (PBS with 0.5% BSA and 0.02% NaN3) and combined into a single 15 mL conical tube. For staining, cells were resuspended in cell-staining media, blocked with FcX (Biolegend, RRID: RRID:AB 1574975) for 5 minutes at room temperature, then incubated for 30 minutes at room temperature on a shaker, with surface marker antibody cocktail (Supplementary Table S5) in 1 mL total volume for tumor, and 500 μL volume for other organs. Cells were washed in cell-staining media then permeabilized with methanol for 10 minutes at 4 °C, then washed again in cell-staining media. Cells were then incubated with the intracellular antibody cocktail in the same volume as surface markers for 30 minutes at room temperature on a shaker. Cells were washed in cell-staining media then incubated with 1 mL PBS containing 1:5000 Iridium intercalator (Fluidigm) and 4% final formaldehyde. Cells were maintained in this final buffer for 2 days prior to sample acquisition on a CyTOF 2 mass cytometer (Fluidigm) using Maxpar Cell Acquisition Solution. 1-5 x 10^5^ cells were acquired per each sample. Data normalization was performed using EQ Four Element Calibration Beads (Fluidigm) included during sample acquisition, using Normalizer v0.3 (Nolan lab). Debarcoding was performed using Single Cell Debarcoder v0.2 (Nolan lab).

### Mass cytometry data analysis

Immune cell subsets were manually gated using the strategy shown in Supplementary Fig. S2. Input cells for DAseq were exported from manually gated cells, with the same number of cells imported per sample. UMAP dimensionality reduction^46^ using the package “uwot” (v.0.1.16) was performed using all available protein markers except Ki-67. Differential abundance analysis (DA analysis) was performed using the package “DAseq” (v.1.0.0) using the top 10 principal components determined from the asinh expression for each marker for each cell. Top DA cells were determined by thresholding cells with positive or negative DA measures beyond the extent of random distribution.

For pairwise correlation analysis involving neutrophil clusters, clusters were first identified using UMAP and Phenograph^108^ using cells from each organ, with manual annotation of clusters representing neutrophils. Cells from LLC MSI-H or B16F10 MSI-H tumor bearing animals were used for further analysis. For each sample, the cluster frequencies were calculated (percent of total singlets for tumor, and percent of live CD45+ cells for other organs). Spearman pairwise correlation was performed as previously described^49^, and filtered for correlations involving neutrophils with r > 0.7 or < −0.7.

For most analyses, batch correction was not applied as aliquots of the same antibody cocktail was used to acquire the entire dataset for analysis. Data in Supplementary Fig. S4 required comparison across experiments acquired separately with antibody cocktails generated at different times. For this analysis, batch correction was applied using the package CyCombine (v. 0.2.16) using an anchor sample common across the two CyTOF runs.

For quantification of immune subsets, cells were quantified as a percentage of total singlets, including live and dead cells, and CD45+ and CD45- cells, to reflect the density of that cell type among cells comprising the harvested organ.

### Cytokine quantification

For quantification of cytokines from tumor, minced tumor was lysed using RIPA buffer containing protease inhibitor (Cell Biolabs) on ice for 10 minutes, followed by centrifugation at 14,000 rpm for 10 minutes at 4 °C. Supernatant containing cell lysate was collected and normalized to the same A595 using the Bio-Rad Protein Assay. Samples were frozen and stored at −80 °C until quantification. For quantification of cytokines from plasma, freshly isolated peripheral blood was centrifuged for 10 minutes at 4 °C at 1000 *g*. 100 μL of supernatant plasma was mixed 1:1 with 100 μL of PBS/EDTA, and frozen until quantification. For multiplex cytokine quantification, lysates or plasma were submitted for mouse discovery assays available through Eve Technologies (Calgary, AB). For G-CSF and CXCL1 ELISA, tumor lysates were analyzed using the mouse G-CSF ELISA Kit (Abcam ab197743) and CXCL1 ELISA Kit (Abcam ab216951) per manufacturer protocol. Principal component analysis was performed using cytokine concentrations as input. Values were log adjusted then analyzed using the function PCA from the R package FactoMineR^109^ (v2.4). Synergy analyses were performed using SynergyFinder Plus web interface (synergyfinder.org^110^), following normalization of G-CSF concentrations with 100% representing the maximum value for each cell line.

### Flow cytometry and cell sorting

All flow cytometry antibodies and concentrations used for analysis is found in Supplementary Table S6. Tissue samples were initially processed as for CyTOF, and cells were stained for viability with Zombie-NIR stain (Biolegend). Cell surface staining was performed in cell-staining media (PBS with 0.5% BSA and 0.02% NaN3) for 20 minutes at room temperature. For intracellular staining, cells were permeabilized and fixed using eBioscience Foxp3/Transcription Factor Staining Buffer Set prior to staining in 1X permeabilization buffer. Stained cells were analyzed with a CytoFLEX flow cytometer (Beckman Coulter) or an LSR II flow cytometer (BD Biosciences). Stained live cells were sorted using FACSAria cell sorters (BD Biosciences) using a 100 μm nozzle and event rate of 3000-4000 per second into collecting vessels kept at 4 °C until downstream use. Gating strategies are found in Supplementary Fig. S9.

### In vitro cellular assays

For tumor-T cell co-culture, CD8 T cells and tumor cells were sorted as above. Prior to plating, 96 well flat bottom plate wells were coated with anti-CD3 antibody (clone 145-2C11, 10 μg/mL final in PBS, UCSF Monoclonal Antibody Core) for 4-6 hours at 37°C and washed twice with PBS. For each cell population in the co-culture, 3.5 x 10^4^ cells were plated together with anti-CD28 antibody (clone 37.51, 3 μg/mL final, UCSF Monoclonal Antibody Core) and any neutralizing antibodies in 200 μL final volume of T cell media (RPMI 1640 with 10% FCS, 25mM HEPES, 2mM L-glutamine, 33μM 2-mercaptoethanol, 100 U mL^-1^ penicillin and 100 μg mL^-1^ streptomycin. After 24 hours of culture in a 37 °C incubator, the supernatant was collected for G-CSF ELISA.

For OT-I T cell MDSC assay, neutrophils were sorted as above from LLC MSI-H tumor, while OT-I cells were harvested from spleen of OT-I CD45.1 mice (RRID:IMSR_JAX:003831) by processing over a 70 μm filter. In 96 well U-bottom plates, 1 x 10^5^ OT-I splenocytes and 2.5 x 10^4^ sorted neutrophils were combined with or without SIINFEKL peptide at 0.4 ng/mL in 200 μL T cell media. Cells were incubated in a 37 °C CO2 incubator for 42 hours, then BrdU was added to the wells at 10 μM final concentration for an additional 6 hours. Cells were collected and stained with Zombie NIR, anti-TCRβ, anti-CD45.1, and anti-CD8a antibodies then anti-BrdU antibody as per manufacturer protocol for the eBioscience BrdU Staining Kit for Flow Cytometry. Cells were analyzed using an LSR II flow cytometer.

For assessment of G-CSF and CXCL1 by ELISA following TNFα, IL-1α, or IL1-β stimulation, 10 ng/ml of cytokine (Biolegend) was added (unless otherwise stated) to cell culture media 24 hours prior to collection of supernatant for analysis using ELISA kits for G-CSF (Abcam, ab197743), or CXCL1 (Abcam, ab216951). For analysis of IκBα by flow cytometry, TNFα and IL-1α (1 ng/ml) was added to culture media 20 minutes prior to cell harvest and processing for staining and flow cytometry as above.

### Quantitative reverse transcription PCR

Cells were sorted into 500 μL TRIzol reagent (Thermo Fisher Scientific). RNA was isolated using Direct- zol 96 MagBead RNA kit (Zymo) per manufacturer protocol. cDNA was generated using Iscript Reverse Transcription Supermix (Bio-Rad) per manufacturer protocol. Real-time PCR was performed using Taqman primers for *Csf3* (Thermo Fisher Scientific Mm00438334_m1), *Cxcl1* (Mm04207460_m1), *Il1a* (Mm00439620_m1), *Il1b* (Mm00434228_m1) and *Gapdh* (Mm99999915_g1) on a Bio-Rad C1000 Thermal Cycler. ΛΛCq values were calculated against *Gapdh* then against biological control. For breast cancer organoid RT-qPCR, Taqman primers for *IL1R1* (Hs00991010_m1), *TNFRSF1A* (Hs01042313_m1), *CXCL8* (Hs00174103_m1), and *UBC* (Hs05002522_g1) were used.

### Western blot

Protein lysates were generated using RIPA buffer containing protease inhibitor (Cell Biolabs) on ice for 10 minutes, followed by centrifugation at 14,000 rpm for 10 minutes at 4 °C. Supernatant containing cell lysate was collected, and 20 μg of lysate was loaded onto NuPAGE 4-12% Bis-Tris protein gel (Thermo Fisher Scientific). Gels were run in NuPAGE MOPS SDS running buffer, then transferred to a nitrocellulose blot in NuPAGE Transfer Buffer. Blots were incubated with PBS-T (0.1% Tween 20) with 5% BSA, then incubated with primary antibody against MSH2 (clone FE11, Thermo Fisher Scientific, 1:500 dilution, RRID: AB_2533139) overnight at 4 °C. After washing in PBST, HRP goat anti-mouse IgG secondary antibody (RRID:AB_562588) was added in PBS-T and incubated for 1 hour. Blot was developed using SuperSignal West Pico PLUS chemiluminescent substrate (Thermo Fisher Scientific) and acquired using a Bio-Rad ChemiDoc system.

### Organoid culture

Organoid cultures were generated from breast tumor samples collected at Brigham & Women’s Hospital and processed on the day of surgery with appropriate informed consent for tissue acquisition for scientific research. Viable tissue was minced and placed in a 50 mL conical tube that contained: 18 mL AdDF+++ (Advanced DMEM/F12 containing Glutamax, HEPES, and antibiotics) + 2 mL collagenase (10 mg/ml). The tube was placed into an orbital shaker at 37 °C for 30 min. After digestion, 30 mL AdDF+++ plus 2% FBS was added to the tube and the pellet was collected, resuspended in BME type 2 (R&D Systems, 353300502), and cultured as previously described^111^. Briefly, a drop of 50 μL of this suspension was placed in the center of a well in a 24-well non- treated culture plate (Corning, 3738) and transferred to a 37°C, 5% CO2 incubator to harden for 20 min.

Subsequently, 500 μL Type 1 breast cancer organoid culture medium^112^ was added to the well. Medium was changed every 2-3 days. TrypLE Express (Gibco, 12604013) was used for passaging approximately every 1–2 weeks. Organoids were viably frozen by Recovery™ Cell Culture Freezing Medium (Gibco, 12648010). TORG40 was generated from a triple-negative inflammatory breast cancer, and TORG139 was generated from a triple-negative breast cancer.

### Human breast cancer single-cell RNAseq analysis

Data was obtained from https://lambrechtslab.sites.vib.be/en/single-cell and is deposited at the European Genome-phenome Archive (EGA) under study no (EGAS00001004809) and under data accession no. EGAD00001006608. Data was imported to Seurat (v. 4.1.0) as a Seurat Object. For quality control, the data was previously filtered by excluding all cells expressing <200 or >6,000 genes, cells that contained <400 unique molecular identifiers (UMIs), and cells with >15% mitochondrial counts. Raw counts were normalized using sctransform (v. 0.3.3) and glmGamPoi (v. 1.2.0) with regression of cell cycle genes, percent mitochrondrial counts, and number of UMIs (counts). Principal Component Analysis (PCA) was performed using variable genes. Cell clusters were defined using Seurat’s FindNeighbors (dim 1-20) and FindClusters (resolution 0.5) and were visualized using Uniform Manifold Approximation and Projection (UMAP). Cluster annotations for major cell types were provided in the metadata and aligned well with cluster assignment. The T cell cluster was fine annotated into CD4, CD8, and Tregs based on expression of *CD4*, *CD8A*, *CD8B*, and *FOXP3*. The data was then filtered to CD8 T cells, tumor cells, CD4 T cells, and myeloid cells.

CD8 T cells were reclustered using the above parameters. Clusters were defined using the FindAllMarkers function in Seurat which identified differentially expressed genes (DEGs). Genes must be expressed in 25% of cells. DEGs between specific clusters, found using FindMarkers, were plotted using EnhancedVolcano (v. 1.8.0). For genes of interest, the density, or for two genes the joint density, was plotted in UMAP space using Nebulosa (v. 1.0.2). DEenrichRPlot (Seurat and enrichR (v. 3.0)) was run to determine GO terms enriched in specific clusters using the GO_Biological_Process_2021 list.

The proportion of cells in a given cluster from “Pre” or “On” treatment was calculated on a per patient basis by normalizing for the number of CD8s per patient. The frequency of any cluster of CD8s compared “Pre” vs “On” treatment was normalized to the number of cells in the entire tumor. Paired differences in cluster frequency were tested using paired Wilcoxon. Tumor cell scoring for NF-κB/TNFα signaling was performed using the gene set “HALLMARK_TNFA_SCORING_VIA_NFKB” obtained from Molecular Signatures Database or MSigDB and the AddModuleScore function in Seurat. Differences in score were calculated “Pre” vs “On” treatment.

### CIBERSORTx analysis

COAD bulk RNA sequencing data from TCGA was analyzed using the CIBERTSORTx^72^ platform (cibersortx.stanford.edu). Proportion of neutrophils as percentage of immune cells in sample was deconvoluted using the LM22 signature matrix and default parameters. MSIsensor^113^ scores for each tumor sample was used to stratify patients as MSS or MSI-H using a published dataset^10^.

### Human peripheral blood neutrophil to lymphocyte ratio analysis

Patients with MSI-H tumors from our institutional database of tumors submitted for next generation sequencing were identified (UCSF IRB 18-24633). Dataset was further filtered to non-central nervous system tumors and patients receiving ICI therapy (included single or combinations of pembrolizumab, nivolumab, atezolizumab, or ipilimumab) during their course of treatment. Among this group of patients, those who experienced radiologist-confirmed imaging-based progression were identified, with the date of progression defined as the date of imaging. 6-week averaged values were derived from all available complete blood count with differential lab testing prior to initiation of immunotherapy or prior to progression of disease. For each patient and timepoint, the absolute neutrophil count was divided by the absolute lymphocyte count to derive the neutrophil to lymphocyte ratio (NLR).

### Human lung cancer dataset analysis

Log2 normalized TPM and relevant clinical data were downloaded from EGA (https://ega-archive.org/studies/EGAS00001005013) for the OAK clinical trial^76^. Single sample gene signature scores were assigned to bulk RNA-seq samples from OAK by taking the median z-score of the genes that comprise the signature (*“IL1R1”,“TNFRSF1A”,“IL1A”,“IL1B”,“TNF”,“CD69”,“CXCL1”,“CXCL2”,“CXCL8”,“NFKB1”,“NFKB2”,“CS*

*F3"*). Patients were grouped into “high” or “low” expression based on the cohort-wide median expression of the signature and also CD8A expression. For resultant survival analyses in the atezolizumab arm of OAK, the log-rank test was used to compare Kaplan-Meier survival curves and cox proportional hazards regression models were used to generate hazard ratios and 95% confidence intervals.

## Statistical analyses

Unless otherwise indicated, statistical testing was performed using GraphPad Prism. Prior to significance testing for differences among groups, the normality of data distribution for each group being tested was tested via Shapiro-Wilk test, where data with p > 0.05 were considered normally distributed. For normal datasets, t test was performed, with Welch’s correction if F test demonstrated unequal SD between groups. For non-normal datasets, Mann-Whitney U test was used. Multiple testing correction was performed using Holm-Sidak method to generate adjusted p values where indicated. Two-tailed tests were performed where direction of change could not be hypothesized a priori, and one tailed-tests were performed to test a specific direction of change. All data points represent biological replicates.

## Data and reagent availability

Mass cytometry data are deposited into Mendeley Data and also available by request to the senior author without restrictions. Bulk RNA sequencing data and exome sequencing data are deposited into NCBI SRA database. All cell lines are available upon request to the senior author without restrictions.

## Code availability

R code for analyses performed within the manuscript will be made available upon request.

## Acknowledgements

We thank the UCSF Flow Cytometry Core and S. Tamaki for CyTOF maintenance. We thank E. Engleman and J. Bluestone for cell lines. We thank A. Marson and T. Roth for CRISPR-Cas9 reagents, protocols and equipment.

We thank the UCSF Clinical Cancer Genomics Laboratory, Molecular Oncology Initiative, and D. Raleigh for access to the UCSF cBioPortal dataset. This study makes use of publicly available data generated by the Lambrechts group in the BioKey study. A full list of investigators who contributed to the generation of the data, and the funding source is available at doi: 10.1038/s41591-021-01323-8/. The Lambrechts group is not responsible for the analysis or interpretation of the data presented. This study was supported by NIH grants DP5 OD023056 and R01 DE032033 to M.H.S., the UCSF Prostate Cancer Program Pilot Award to N.W.C., and NIH S10 OD018040 to procure the CyTOF mass cytometer. N.W.C. is supported by NIH grant T32 5T32AI007334- 33, and ASTRO/ACS Clinician Scientist Development Grant ASTRO-CSDG-23-1036400-01-IBCD. S.M.G. is supported by NIH grants F31CA271748, T32GM1365471, and T32GM00856825, and the UCSF ImmunoX Computational Biology Initiative. K.J.H-G is supported by the Stanford Propel Postdoctoral Scholarship. R.D. is supported by NIH grant F31 CA265128. J.L.Y. is supported by an NSF GRFP fellowship. M.H.S. is an Investigator of the Parker Institute for Cancer Immunotherapy and of the Chan Zuckerberg Biohub. Breast organoid work by J.Z.Y., F.L., D.A.D., and J.M.R. was supported by METAvivor and the Dana-Farber/Harvard Cancer Center Breast SPORE 1P50CA168504.

## Contributions

Conceptualization: N.W.C., S.M.G., and M.H.S.; experimental methodology: N.W.C., K.J.H-G., R.D., J.L.Y.,

N.A.A., I.T., J.Z.Y., F.L., D.A.D., J.M.R.; computational methodology: N.W.C., S.M.G., E.J.K., K.J.H-G.,

B.Y.N., M.H.S., investigation: all authors; writing, original draft: N.W.C., S.M.G.; writing, review and editing: all authors; funding acquisition: M.H.S. and N.W.C.; supervision: M.H.S.

