## Supplementary Data for "T cells Instruct Immune Checkpoint Inhibitor Therapy Resistance in Tumors Responsive to IL-1 and TNFα Inflammation"

### Figure S1

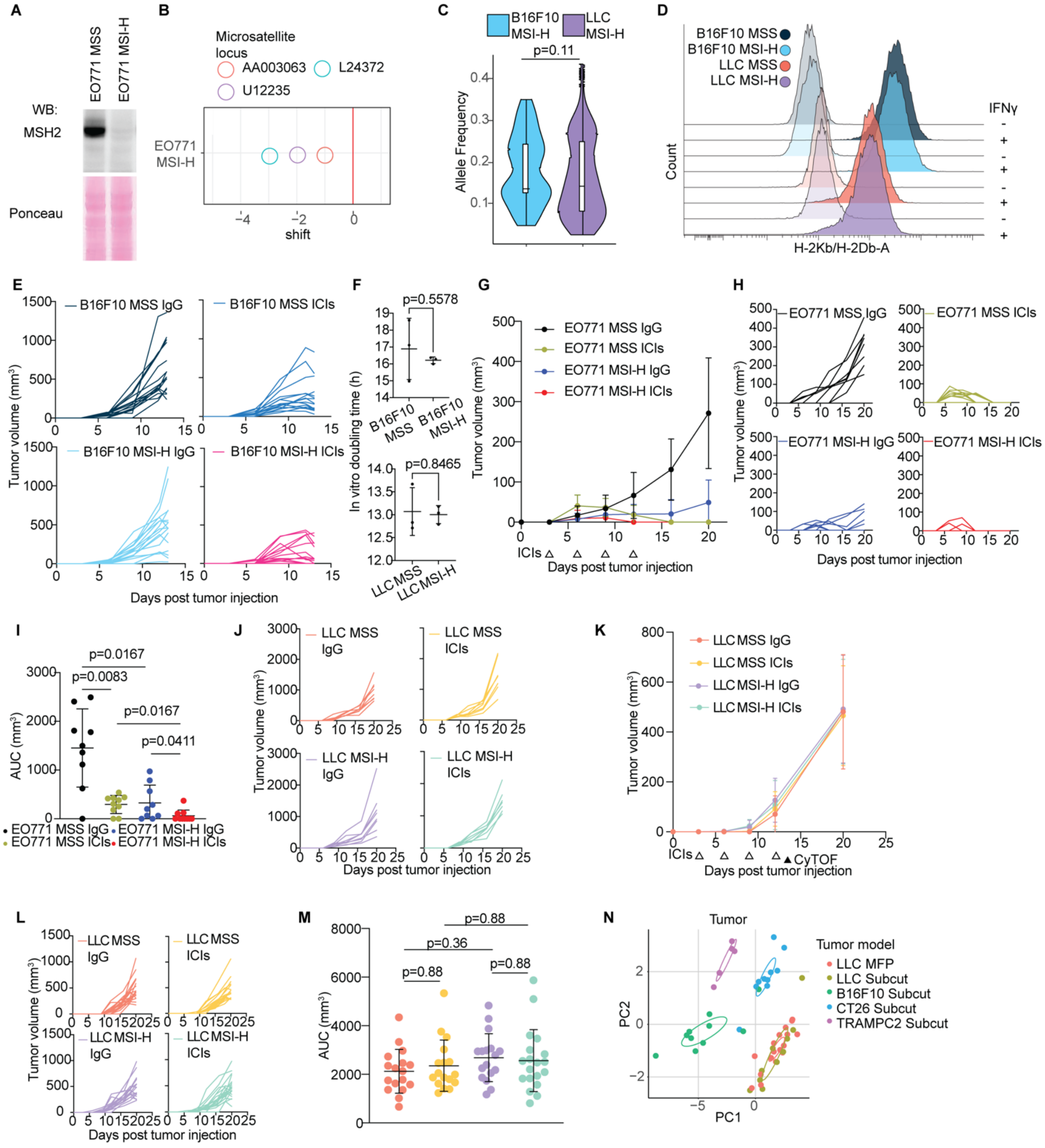

A
EO771 MSS
EO771 MSI-H
WB:
MSH2
Ponceau
B
Microsatellite locus
AA003063
L24372
U12235
EO771 MSI-H
shift
C
B16F10 MSI-H
LLC MSI-H
p=0.11
Allele Frequency
D
B16F10 MSS
B16F10 MSI-H
LLC MSS
LLC MSI-H
IFNγ
Count
H-2Kb/H-2Db-A
E
B16F10 MSS IgG
B16F10 MSS ICIs
B16F10 MSI-H IgG
B16F10 MSI-H ICIs
Tumor volume (mm³)
Days post tumor injection
F
p=0.5578
B16F10 MSS
B16F10 MSI-H
p=0.8465
LLC MSS
LLC MSI-H
In vitro doubling time (h)
G
EO771 MSS IgG
EO771 MSS ICIs
EO771 MSI-H IgG
EO771 MSI-H ICIs
Tumor volume (mm³)
ICIs
Days post tumor injection
H
EO771 MSS IgG
EO771 MSS ICIs
EO771 MSI-H IgG
EO771 MSI-H ICIs
Tumor volume (mm³)
Days post tumor injection
I
p=0.0167
p=0.0083
p=0.0167
p=0.0411
AUC (mm³)
EO771 MSS IgG
EO771 MSI-H IgG
EO771 MSS ICIs
EO771 MSI-H ICIs
J
LLC MSS IgG
LLC MSS ICIs
LLC MSI-H IgG
LLC MSI-H ICIs
Tumor volume (mm³)
Days post tumor injection
K
LLC MSS IgG
LLC MSS ICIs
LLC MSI-H IgG
LLC MSI-H ICIs
Tumor volume (mm³)
ICIs
CyTOF
Days post tumor injection
L
LLC MSS IgG
LLC MSS ICIs
LLC MSI-H IgG
LLC MSI-H ICIs
Tumor volume (mm³)
Days post tumor injection
M
p=0.88
p=0.36
p=0.88
p=0.88
AUC (mm³)
N
Tumor
Tumor model
LLC MFP
LLC Subcut
B16F10 Subcut
CT26 Subcut
TRAMPC2 Subcut
PC1
PC2

**Supplementary Figure S1.**

**A,** Western blot of MSH2 protein in EO771 MSS vs. MSI-H cell line whole cell lysates. Representative of two independent experiments. **B,** Targeted microsatellite PCR assay comparing amplicon size at indicated loci for the EO771 MSI-H vs. MSS cell line. Representative of two independent experiments. **C,** Box and violin plots of neoepitope allele frequencies in indicated cell lines. Two-tailed p value by Mann-Whitney test. Box plot: center line, median; box limits, upper and lower quartiles; whiskers, 1.5x interquartile range. **D,** Quantification of H-2Kb or H-2Db expression by flow cytometry of indicated cell lines *in vitro* and treated with or without IFNγ (62.5 ng/mL for 48 hours). Representative plot from two independent experiments with n=3 each. **E,** Growth curves for individual mice shown in **Fig. 1F and G. F,** Cell doubling times *in vitro* for the indicated cell lines, n=3, representative of two independent experiments. Two-tailed p values by t test. **G and H,** Quantification of mean and individual mouse volumes for EO771 MSS or MSI-H tumors treated with or without anti-PD1 and anti-CTLA4 antibodies (ICIs) at the indicated timepoints (arrowheads). n=9-10 mice per condition from two independent experiments. **I,** Quantification of area under curve (AUC) for the experiment shown in **G and H. J,** Growth curves for individual mice with subcutaneous LLC tumors shown in **Fig. 1H and I. K,** Quantification of mean volumes and area under curves (AUC) for LLC mammary fat pad tumors treated with or without anti-PD1 and anti-CTLA4 antibodies (ICIs) at the indicated timepoints (white arrowhead). Black arrowhead denotes timepoint for sample collection for CyTOF analysis. n=17-19 mice per condition from four independent experiments. Two-tailed adjusted p value by Mann-Whitney test. **L,** Growth curves for individual mice shown in **(K). M,** Quantification of AUC for the experiment shown in **(K). N,** Principal component analysis of manually gated intratumoral immune cell frequencies as percent of singlets in tumor for the indicated tumor types and site of implantation. MFP, mammary fat pad. Subcut, subctutaneous. 95% confidence ellipses are shown. Two-tailed p value by Mann-Whitney tests. For all applicable panels, error bars represent mean +/- SD.

#### Figure S2

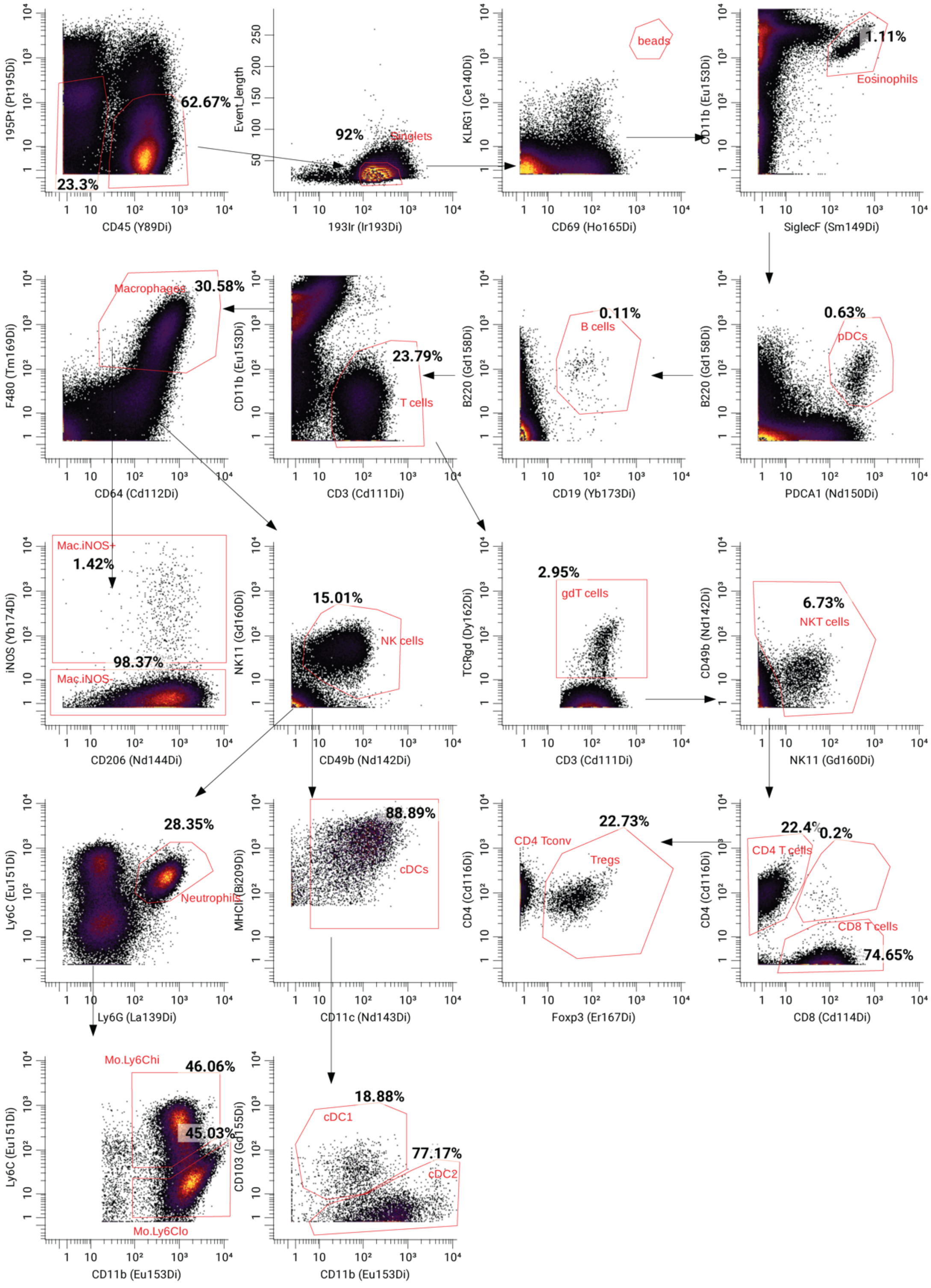
195Pt (Pt195Di)
CD45 (Y89Di)
62.67%
23.3%
Event_length
193Ir (Ir193Di)
92%
Singlets
KLRG1 (Ce140Di)
CD69 (Ho165Di)
beads
CD11b (Eu153Di)
SiglecF (Sm149Di)
1.11%
Eosinophils
F480 (Tm169Di)
CD64 (Cd112Di)
Macrophages
30.58%
CD11b (Eu153Di)
CD3 (Cd111Di)
23.79%
T cells
B220 (Gd158Di)
CD19 (Yb173Di)
0.11%
B cells
B220 (Gd158Di)
PDCA1 (Nd150Di)
0.63%
pDCs
iNOS (Yb174Di)
CD206 (Nd144Di)
Mac.iNOS+
1.42%
98.37%
Mac.iNOS-
NK11 (Gd160Di)
CD49b (Nd142Di)
15.01%
NK cells
TCRgd (Dy162Di)
CD3 (Cd111Di)
2.95%
gdT cells
CD49b (Nd142Di)
NK11 (Gd160Di)
6.73%
NKT cells
Ly6C (Eu151Di)
Ly6G (La139Di)
28.35%
Neutrophils
MHCII (Bi209Di)
CD11c (Nd143Di)
88.89%
cDCs
CD4 (Cd116Di)
Foxp3 (Er167Di)
CD4 Tconv
22.73%
Tregs
CD4 (Cd116Di)
CD8 (Cd114Di)
22.4%
0.2%
CD4 T cells
CD8 T cells
74.65%
Ly6C (Eu151Di)
CD11b (Eu153Di)
Mo.Ly6Chi
46.06%
45.03%
Mo.Ly6Clo
CD103 (Gd155Di)
CD11b (Eu153Di)
18.88%
cDC1
77.17%
cDC2

**Supplementary Figure S2.**

Biaxial plots showing immune subset gating from CyTOF data.

#### Figure S3

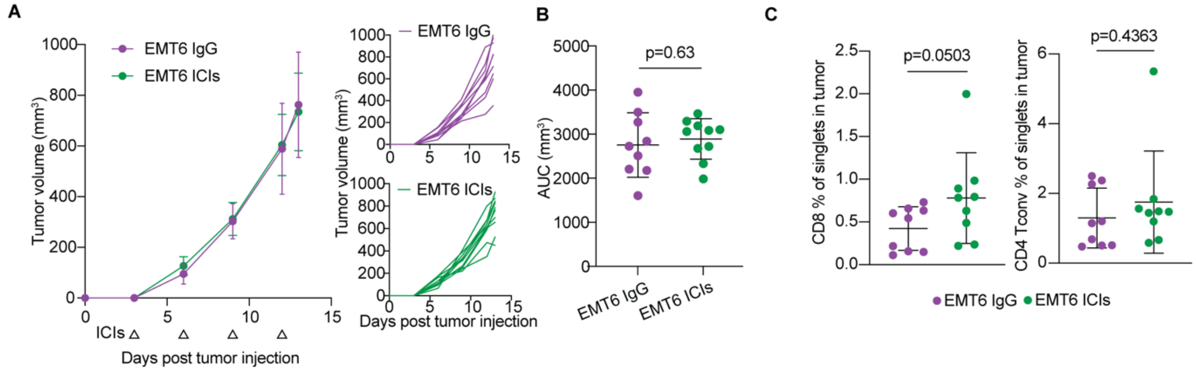

A
EMT6 IgG
EMT6 ICIs
Tumor volume (mm³)
ICIs
Days post tumor injection
B
p=0.63
AUC (mm³)
EMT6 IgG
EMT6 ICIs
C
p=0.0503
CD8 % of singlets in tumor
p=0.4363
CD4 Tconv % of singlets in tumor
EMT6 IgG
EMT6 ICIs


27

**Supplementary Figure S3.**

**A,** Quantification of mean and individual mouse volumes for EMT6 tumors treated with or without anti-PD1 and anti-CTLA4 antibodies (ICIs) at the indicated timepoints (arrowheads). n=9-10 mice per condition from two independent experiments. **B,** Quantification of AUC for the experiment shown in **(A).** Two-tailed p value by t test. **C,** Quantification of CD8 (left panel) and CD4 Tconv (right panel) infiltration in EMT6 tumors at day 13 post implantation from two independent experiments.

**Figure S4**

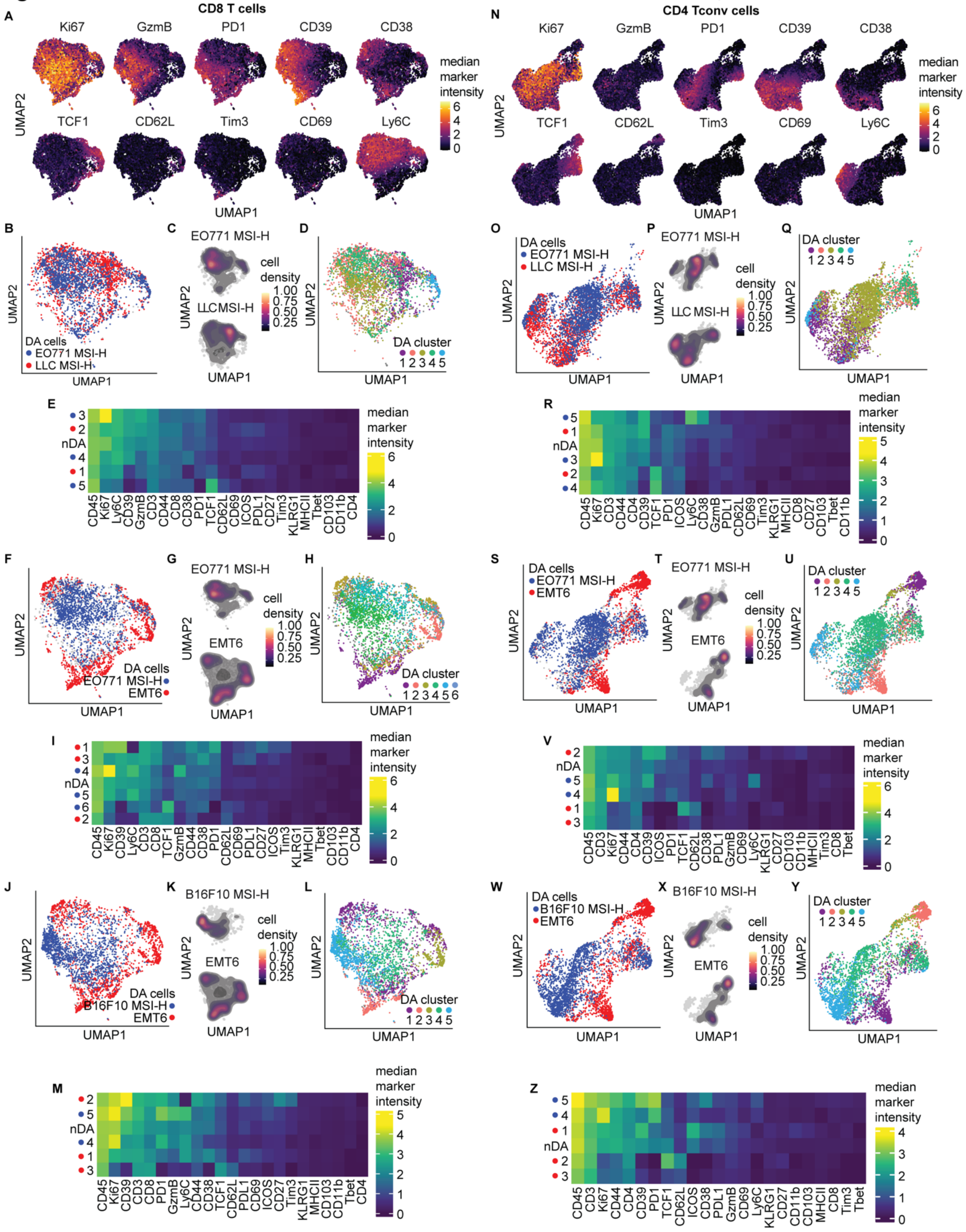
ICIs
CD8 T cells
ICIs
CD4 Tconv cells
Ki67
GzmB
PD1
CD39
CD38
TCF1
CD62L
Tim3
CD69
Ly6C
median marker intensity
UMAP1
UMAP2
DA cells
EO771 MSI-H
LLC MSI-H
EMT6
B16F10 MSI-H
cell density
DA cluster
nDA

**Supplementary Figure S4.**

**A,** Feature plots of selected markers colored by asinh transformed staining intensity from 7,546 CD44+ CD8 T cells from ICI-treated B16F10 MSI-H, LLC MSI-H, EMT6 (day 13), and EO771 MSI-H (day 10) from n=5-6 mice per condition. **B,** Differential abundance (DA) analysis of CD44+ CD8 T cells from EO771 MSI-H and LLC MSI-H tumors. **C,** Density plots for cells in **(B). D,** Clustering of DA cells from **(B)** into unsupervised cell subsets. **E,** Heatmap of median marker intensities for cells in each DA cluster. nDA, non-DA cells. Colored dots indicate clusters more abundant in EO771 (blue) or LLC (red). **F-I,** DA analysis as in **(B-E)** for EO771 MSI-H vs. EMT6 CD44+ CD8 T cells. **J-M,** DA analysis as in **(B-E)** for B16F10 MSI-H vs. EMT6 CD44+ CD8 T cells. **N-Z,** Feature plots and DA analysis as performed in **(A-M)** for 9,366 CD44+ CD4 Tconv cells.

#### Figure S5

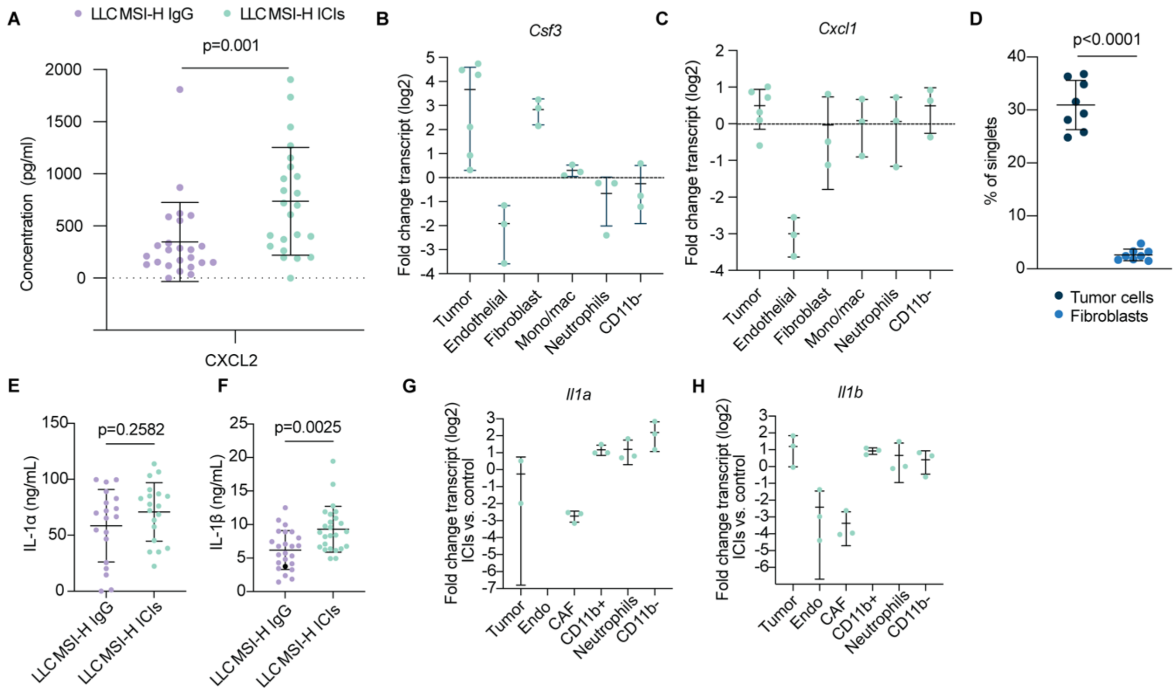

A
LLC MSI-H IgG
LLC MSI-H ICIs
p=0.001
Concentration (pg/ml)
CXCL2
B
Csf3
Fold change transcript (log2)
Tumor
Endothelial
Fibroblast
Mono/mac
Neutrophils
CD11b-
C
Cxcl1
Fold change transcript (log2)
D
p<0.0001
% of singlets
Tumor cells
Fibroblasts
E
p=0.2582
IL-1α (ng/mL)
LLC MSI-H IgG
LLC MSI-H ICIs
F
p=0.0025
IL-1β (ng/mL)
G
Il1a
Fold change transcript (log2) ICIs vs. control
Tumor
Endo
CAF
CD11b+
Neutrophils
CD11b-
H
Il1b


46

**Supplementary Figure S5.**

**A,** Quantification of CXCL2 in day 20 tumor lysates by multiplexed bead assay, from 5 independent experiments. Two-tailed p value by Mann-Whitney test. **B,** *Csf3* transcript quantification from ICI-treated LLC MSI-H tumors as fold change from control, corresponding to experiment shown in **Fig. 3F. C,** *Cxcl1* transcript quantification as in **(B). D,** Quantification of LLC tumor cell and fibroblasts as percentage of singlet cells in tumor, n=8 mice from one experiment. P value by two-tailed Welch's t test. **E,** Quantification of IL-1α from day 20 tumor lysates by multiplexed bead assay in 5 independent experiments. Two-tailed p values by Mann-Whitney test. **F,** Quantification of IL-1β as in **(E). G,** *Il1a* transcript quantification from ICI-treated LLC MSI-H tumors as fold change from control, from one experiment with three biological replicates. **H,** *Il1b* transcript quantification as in **(G).** For all applicable panels, error bars represent mean +/- SD.

**Figure S6**

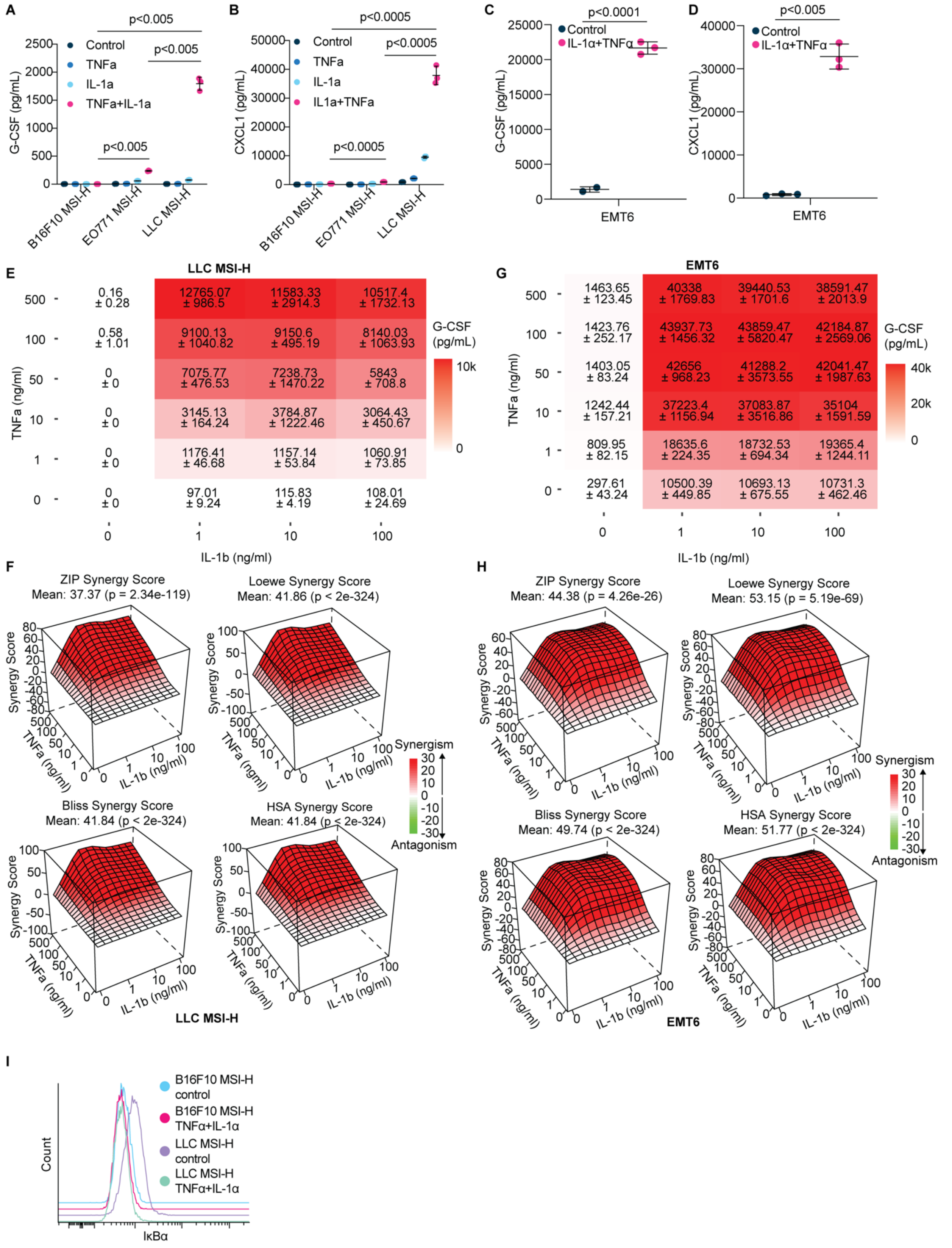

A
p<0.005
p<0.005
p<0.005
Control
TNFa
IL-1a
TNFa+IL-1a
G-CSF (pg/mL)
B16F10 MSI-H
EO771 MSI-H
LLC MSI-H
B
p<0.0005
p<0.0005
p<0.0005
Control
TNFa
IL-1a
IL1a+TNFa
CXCL1 (pg/mL)
B16F10 MSI-H
EO771 MSI-H
LLC MSI-H
C
p<0.0001
Control
IL-1α+TNFα
G-CSF (pg/mL)
EMT6
D
p<0.005
Control
IL-1α+TNFα
CXCL1 (pg/mL)
EMT6
E
LLC MSI-H
TNFa (ng/ml)
500: 0.16 ± 0.28; 12765.07 ± 986.5; 11583.33 ± 2914.3; 10517.4 ± 1732.13
100: 0.58 ± 1.01; 9100.13 ± 1040.82; 9150.6 ± 495.19; 8140.03 ± 1063.93
50: 0 ± 0; 7075.77 ± 476.53; 7238.73 ± 1470.22; 5843 ± 708.8
10: 0 ± 0; 3145.13 ± 164.24; 3784.87 ± 1222.46; 3064.43 ± 450.67
1: 0 ± 0; 1176.41 ± 46.68; 1157.14 ± 53.84; 1060.91 ± 73.85
0: 0 ± 0; 97.01 ± 9.24; 115.83 ± 4.19; 108.01 ± 24.69
IL-1b (ng/ml): 0, 1, 10, 100
G-CSF (pg/mL)
10k
0
G
EMT6
TNFa (ng/ml)
500: 1463.65 ± 123.45; 40338 ± 1769.83; 39440.53 ± 1701.6; 38591.47 ± 2013.9
100: 1423.76 ± 252.17; 43937.73 ± 1456.32; 43859.47 ± 5820.47; 42184.87 ± 2569.06
50: 1403.05 ± 83.24; 42656 ± 968.23; 41288.2 ± 3573.55; 42041.47 ± 1987.63
10: 1242.44 ± 157.21; 37223.4 ± 1156.94; 37083.87 ± 3516.86; 35104 ± 1591.59
1: 809.95 ± 82.15; 18635.6 ± 224.35; 18732.53 ± 694.34; 19365.4 ± 1244.11
0: 297.61 ± 43.24; 10500.39 ± 449.85; 10693.13 ± 675.55; 10731.3 ± 462.46
IL-1b (ng/ml): 0, 1, 10, 100
G-CSF (pg/mL)
40k
20k
0
F
ZIP Synergy Score
Mean: 37.37 (p = 2.34e-119)
Loewe Synergy Score
Mean: 41.86 (p < 2e-324)
Bliss Synergy Score
Mean: 41.84 (p < 2e-324)
HSA Synergy Score
Mean: 41.84 (p < 2e-324)
Synergy Score
TNFa (ng/ml)
IL-1b (ng/ml)
Synergism
Antagonism
LLC MSI-H
H
ZIP Synergy Score
Mean: 44.38 (p = 4.26e-26)
Loewe Synergy Score
Mean: 53.15 (p = 5.19e-69)
Bliss Synergy Score
Mean: 49.74 (p < 2e-324)
HSA Synergy Score
Mean: 51.77 (p < 2e-324)
Synergy Score
TNFa (ng/ml)
IL-1b (ng/ml)
Synergism
Antagonism
EMT6
I
B16F10 MSI-H control
B16F10 MSI-H TNFα+IL-1α
LLC MSI-H control
LLC MSI-H TNFα+IL-1α
Count
IκBα

**Supplementary Figure S6.**

**A,** Dot plot of G-CSF ELISA quantification values from supernatants of indicated cell lines incubated with indicated cytokines for 24h. n=3 biological replicates from a representative experiment, of two independent experiments. P values by two-tailed adjusted Welch's t tests. **B,** Analysis as in **(A)** for CXCL1. **C and D,** Analysis as in **(A)** and **(B)** for EMT6. **E,** G-CSF ELISA quantification from supernatants of LLC MSI-H cells treated with indicated concentrations of IL-1b and TNFa. Values represent mean +/- SD from three biological replicates. **F,** Synergy scores using four synergy models. Mean value of synergy score across the matrix is shown, with p value evaluating the null hypothesis of no interaction between IL-1b and TNFa. **G and H,** Analyses as in **(E)** and **(F)** for EMT6. **I,** Representative histogram of IκBa fluorescence staining intensity in indicated cell lines and treatment conditions.

**Figure S7**

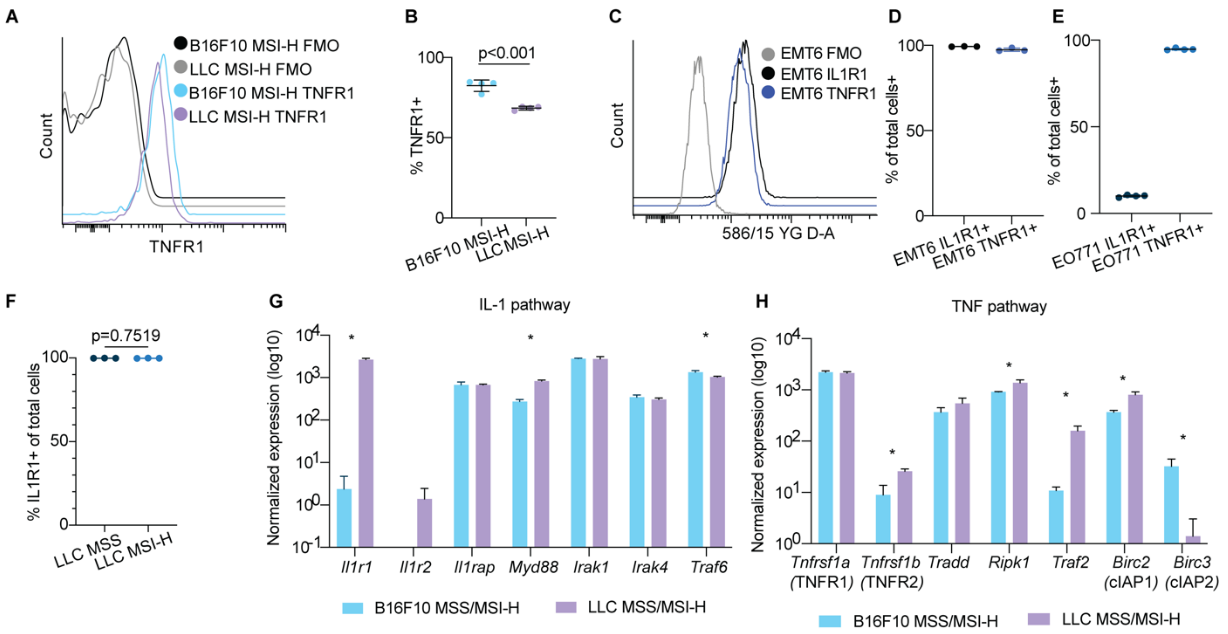
A
B16F10 MSI-H FMO
LLC MSI-H FMO
B16F10 MSI-H TNFR1
LLC MSI-H TNFR1
Count
TNFR1
B
p<0.001
% TNFR1+
B16F10 MSI-H
LLC MSI-H
C
EMT6 FMO
EMT6 IL1R1
EMT6 TNFR1
Count
586/15 YG D-A
D
% of total cells+
EMT6 IL1R1+
EMT6 TNFR1+
E
% of total cells+
EO771 IL1R1+
EO771 TNFR1+
F
p=0.7519
% IL1R1+ of total cells
LLC MSS
LLC MSI-H
G
IL-1 pathway
Normalized expression (log10)
Il1r1
Il1r2
Il1rap
Myd88
Irak1
Irak4
Traf6
B16F10 MSS/MSI-H
LLC MSS/MSI-H
H
TNF pathway
Normalized expression (log10)
Tnfrsf1a (TNFR1)
Tnfrsf1b (TNFR2)
Tradd
Ripk1
Traf2
Birc2 (cIAP1)
Birc3 (cIAP2)
B16F10 MSS/MSI-H
LLC MSS/MSI-H

70

**Supplementary Figure S7.**

**A,** Representative histogram of TNFR1 fluorescence staining intensity for indicated cell lines. **B,** Quantification of % TNFR1+ population in indicated cell lines, from one experiment with n=4 biological replicates. Two independent experiments were performed. P value by two-tailed t test. **C,** Representative histogram of IL1R1 and TNFR1 fluorescence staining intensity for EMT6 cell line. **D,** Quantification of % IL1R1+ and % TNFR1+ populations for EMT6 cells, n=3 biological replicates from one representative experiment. Two independent experiments were performed. **E,** Quantification of % IL1R1+ and % TNFR1+ populations for EO771 MSI-H cells, n=4 biological replicates from one representative experiment. Two independent experiments were performed. **F,** Quantification of % IL1R1+ populations for LLC MSS vs. LLC MSI-H cells. n=3 biological replicates from one representative experiment. p value by t test. Two independent experiments were performed. **G,** Quantification of normalized transcript counts for indicated genes belonging to the IL-1 receptor signaling pathway for the indicated cell lines, n=4 biological replicates per condition. Asterisks indicate $p<0.05$ by Mann-Whitney test. **H,** Quantification as in **(G)** for genes in the TNF receptor signaling pathway. For all applicable panels, error bars represent mean +/- SD.

#### Figure S8

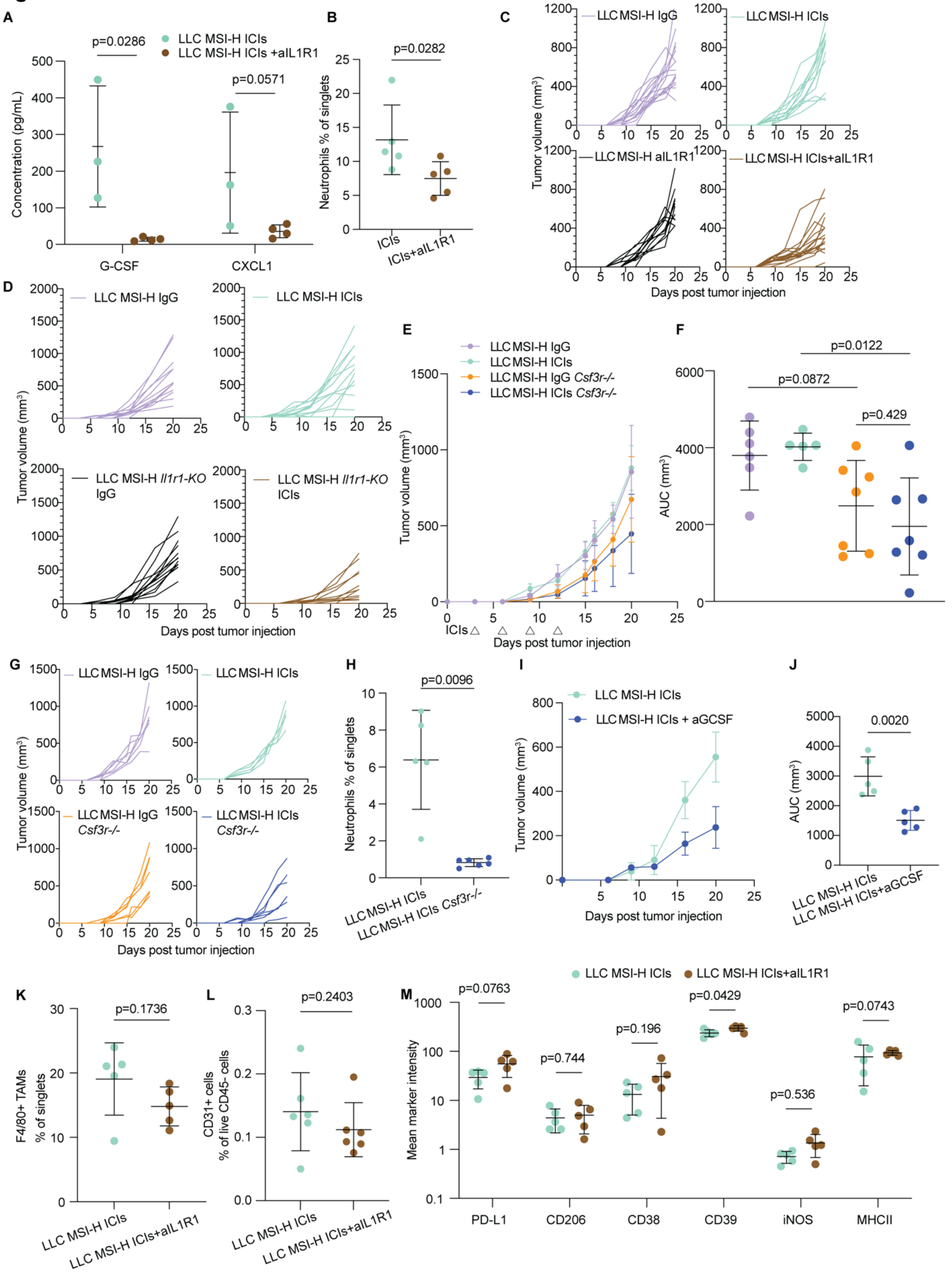

A
p=0.0286
p=0.0571
LLC MSI-H ICIs
LLC MSI-H ICIs +aIL1R1
Concentration (pg/mL)
G-CSF
CXCL1
B
p=0.0282
Neutrophils % of singlets
ICIs
ICIs+aIL1R1
C
LLC MSI-H IgG
LLC MSI-H ICIs
LLC MSI-H aIL1R1
LLC MSI-H ICIs+aIL1R1
Tumor volume (mm³)
Days post tumor injection
D
LLC MSI-H IgG
LLC MSI-H ICIs
LLC MSI-H Il1r1-KO IgG
LLC MSI-H Il1r1-KO ICIs
Tumor volume (mm³)
Days post tumor injection
E
LLC MSI-H IgG
LLC MSI-H ICIs
LLC MSI-H IgG Csf3r-/-
LLC MSI-H ICIs Csf3r-/-
Tumor volume (mm³)
ICIs
Days post tumor injection
F
p=0.0122
p=0.0872
p=0.429
AUC (mm³)
G
LLC MSI-H IgG
LLC MSI-H ICIs
LLC MSI-H IgG Csf3r-/-
LLC MSI-H ICIs Csf3r-/-
Tumor volume (mm³)
Days post tumor injection
H
p=0.0096
Neutrophils % of singlets
LLC MSI-H ICIs
LLC MSI-H ICIs Csf3r-/-
I
LLC MSI-H ICIs
LLC MSI-H ICIs + aGCSF
Tumor volume (mm³)
Days post tumor injection
J
0.0020
AUC (mm³)
LLC MSI-H ICIs
LLC MSI-H ICIs+aGCSF
K
p=0.1736
F4/80+ TAMs % of singlets
LLC MSI-H ICIs
LLC MSI-H ICIs+aIL1R1
L
p=0.2403
CD31+ cells % of live CD45- cells
LLC MSI-H ICIs
LLC MSI-H ICIs+aIL1R1
M
LLC MSI-H ICIs
LLC MSI-H ICIs+aIL1R1
p=0.0763
p=0.744
p=0.196
p=0.0429
p=0.536
p=0.0743
Mean marker intensity
PD-L1
CD206
CD38
CD39
iNOS
MHCII


85

**Supplementary Figure S8**

**A,** Quantification of G-CSF and CXCL1 in day 20 tumor lysates following indicated *in vivo* treatments, n=3-4 mice from one experiment. P values by one-tailed Mann-Whitney test. **B,** Quantification of neutrophils in LLC tumors treated with indicated antibodies and harvested at day 13, from one experiment with n=5 mice. P value by one-tailed t test. **C,** Growth curves for individual mice shown in **Fig. 4A. D,** Growth curves for individual mice shown in **Fig. 4C. E and F,** Quantification of mean volumes and area under curve (AUC) for LLC MSI-H tumors implanted into wild type vs. *Csf3r-/-* G-CSF Receptor knockout mice. n=5-7 mice per condition from one experiment. Two-tailed adjusted p value by t test. **G,** Individual growth curves for mice in experiment shown in **(E). H,** Quantification of neutrophils as percent of singlets in tumor from day 20 LLC MSI-H tumors treated with ICIs and implanted into wild type vs. *Csf3r-/-* mice. n=5-6 per condition from one experiment. P value by two-tailed t test. **I and J,** Quantification of mean volumes and AUC for LLC MSI-H tumors with or without G-CSF neutralizing antibody treatment (aGCSF). n=5 from one representative *in vivo* experiment from three independent experiments. P value by two-tailed t test. **K,** Quantification of F4/80+ tumor-associated macrophages (TAMs) from day 20 LLC MSI-H tumors treated with or without IL1R1 neutralizing antibody. n=5 mice from one experiment. p value by two-tailed t test. **L,** Quantification of CD31+ endothelial cells as for **(K). M,** Quantification of mean marker intensity for indicated markers for F4/80+ TAMs in LLC MSI-H ICI treated tumors with or without aIL1R1 treatment. n=5 mice from one experiment. P value by two-tailed t tests. For all applicable panels, error bars represent mean +/- SD.

#### Figure S9

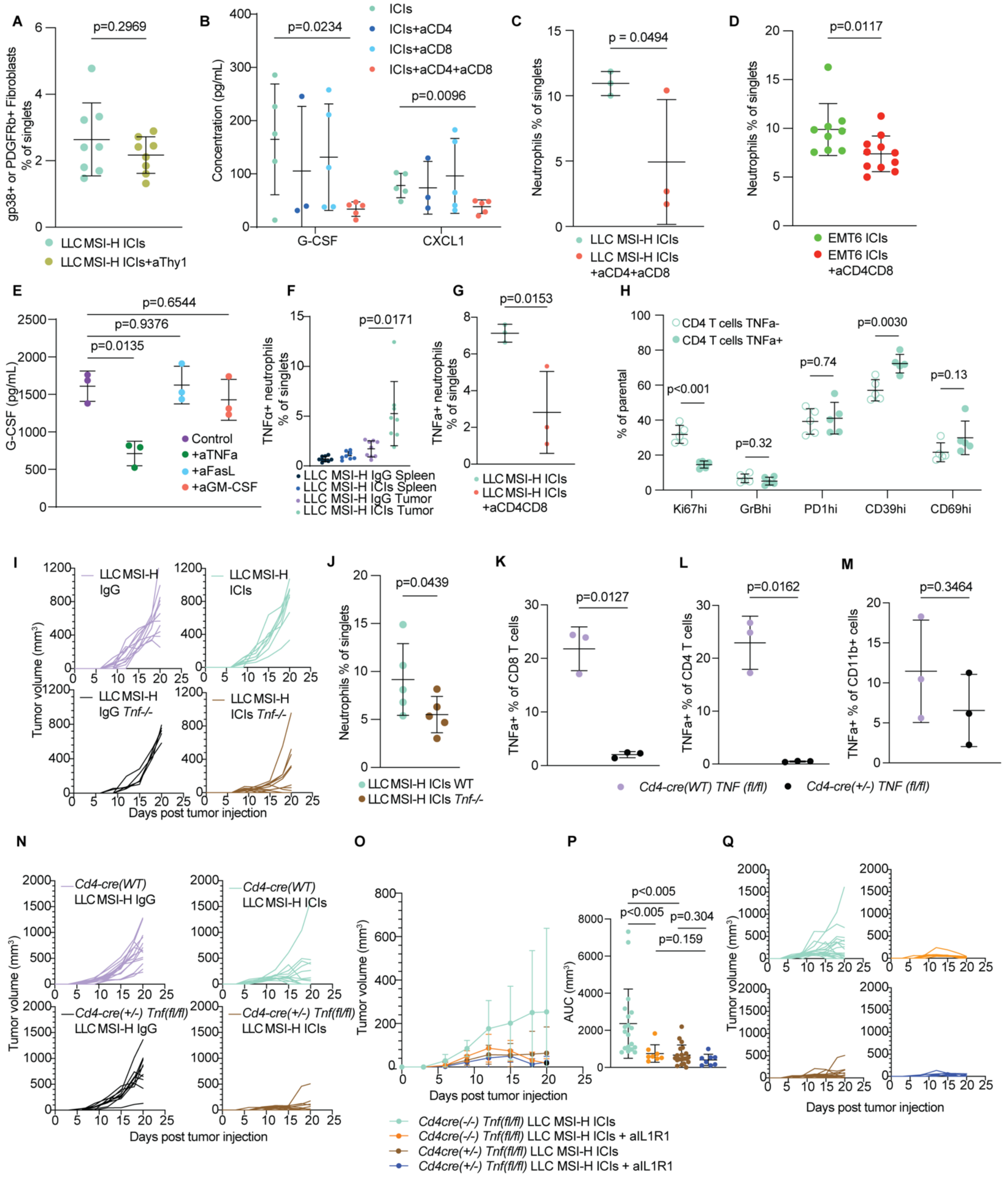


104

**Supplementary Figure S9**

**A,** Quantification of fibroblasts in tumor harvested at day 20 following treatment with indicated antibodies, n=8 mice from one experiment. P value by two-tailed t test. **B,** Quantification of G-CSF and CXCL1 concentration from day 20 tumor lysates for LLC MSI-H tumors treated with the indicated antibodies, n=3-5 mice from one experiment. Adjusted p values by two-tailed Welch's t test. **C,** Quantification of neutrophils in LLC MSI-H tumors, n=3 mice from one experiment. P value by one-tailed t test. **D,** Quantification of neutrophils in EMT6 tumors at day 13 post implantation, n=9-11 mice from two independent experiments. P value by one-tailed t test. **E,** Quantification of G-CSF concentration in supernatant of 24h culture containing indicated sorted cell types from LLC MSI-H tumors treated with ICIs, with or without addition of indicated neutralizing antibodies. Adjusted p values from t tests. n=3 mice from one experiment. **F,** Quantification of TNFα+ neutrophils in tumor or spleen of day 20 LLC MSI-H tumors treated with indicated antibodies, n=8-9 mice from two independent experiments. P value by two-tailed Welch's t test. **G,** Quantification of TNFα+ neutrophils in day 20 tumors following treatment with the indicated antibodies, n=3 mice from one experiment. P value by one-tailed t test. **H,** Quantification of TNFα- and TNFα+ CD8 T cells as % of cells binned into high expression gates for each marker. n=5 mice from one representative experiment. Two independent experiments were performed. P value by two-tailed t tests. **I,** Growth curves for individual mice shown in **Fig. 5I and J. J,** Quantification of neutrophils in LLC MSI-H tumors grown in mice with indicated genotypes, at day 13 post implantation, n=5 mice from one experiment. P value by one-tailed t test. **K,** Quantification of TNFα+ CD8 T cells among stimulated splenocytes from mice with indicated genotypes, from one experiment with n=3 mice. P value from two-tailed Welch's t test. **L,** Quantification as in **(K)** for CD4 T cells. **M,** Quantification as in **(K)** for CD11b+ myeloid cells. **N,** Growth curves for individual mice shown in **Fig. 5K and L. O and P,** Quantification of mean volumes and AUCs for ICI-treated LLC MSI-H tumors implanted into mice with indicated genotypes and treated with or without aIL1R1 treatment. n=8-20 mice from 3 independent experiments. Adjusted p value by Mann-Whitney tests. **Q,** Growth curves for individual mice shown in **(O).**

**Figure S10**

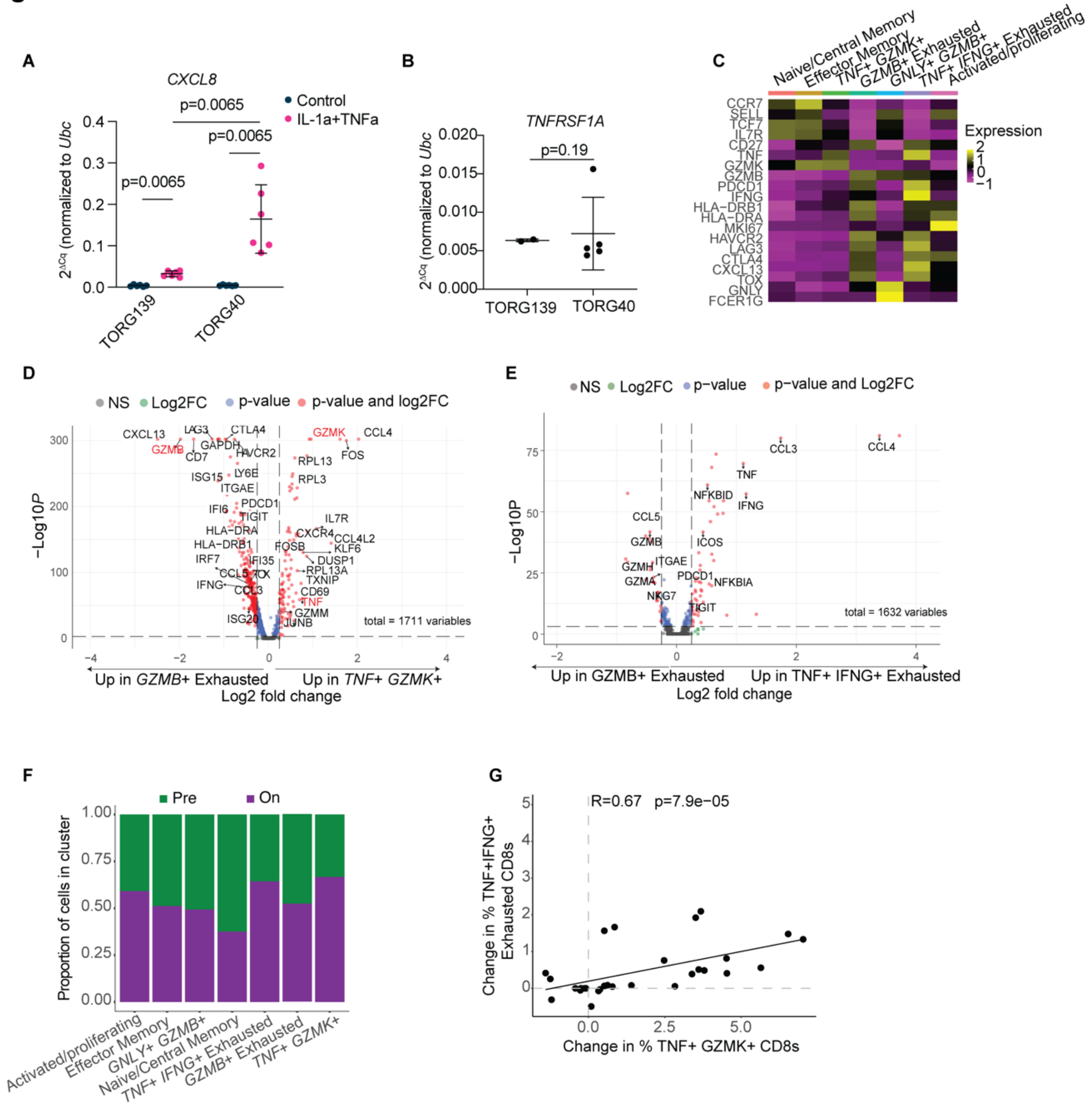


129

**Supplementary Figure S10**

**A,** *CXCL8* transcript quantification by RT-qPCR for breast organoid models, n=6 biological replicates from two independent experiments. **B,** *TNFR1* transcript quantification by RT-qPCR for breast organoid models from two independent experiments. P value by one-tailed Mann-Whitney test. **C,** Heatmap of genes defining CD8 T cell clusters. **D,** Volcano plot of differential gene expression between *TNF*+ *GZMK*+ Exhausted CD8s and *GZMB*+ CD8 T cells. **E,** Volcano plot of differential gene expression between *TNF*+ *IFNG*+ CD8s and *GZMB*+ Exhausted CD8 T cells. **F,** Stacked bar plot of proportion of cells in each cluster originating from pre-treatment samples (Pre) vs. on-treatment (On). Proportions are normalized for number of CD8 T cells per patient. **G,** Spearman correlation analysis between the change in % of *TNF*+ *IFNG*+ CD8 T cells and the change in *TNF*+*GZMK*+ CD8 T cells.

#### Figure S11

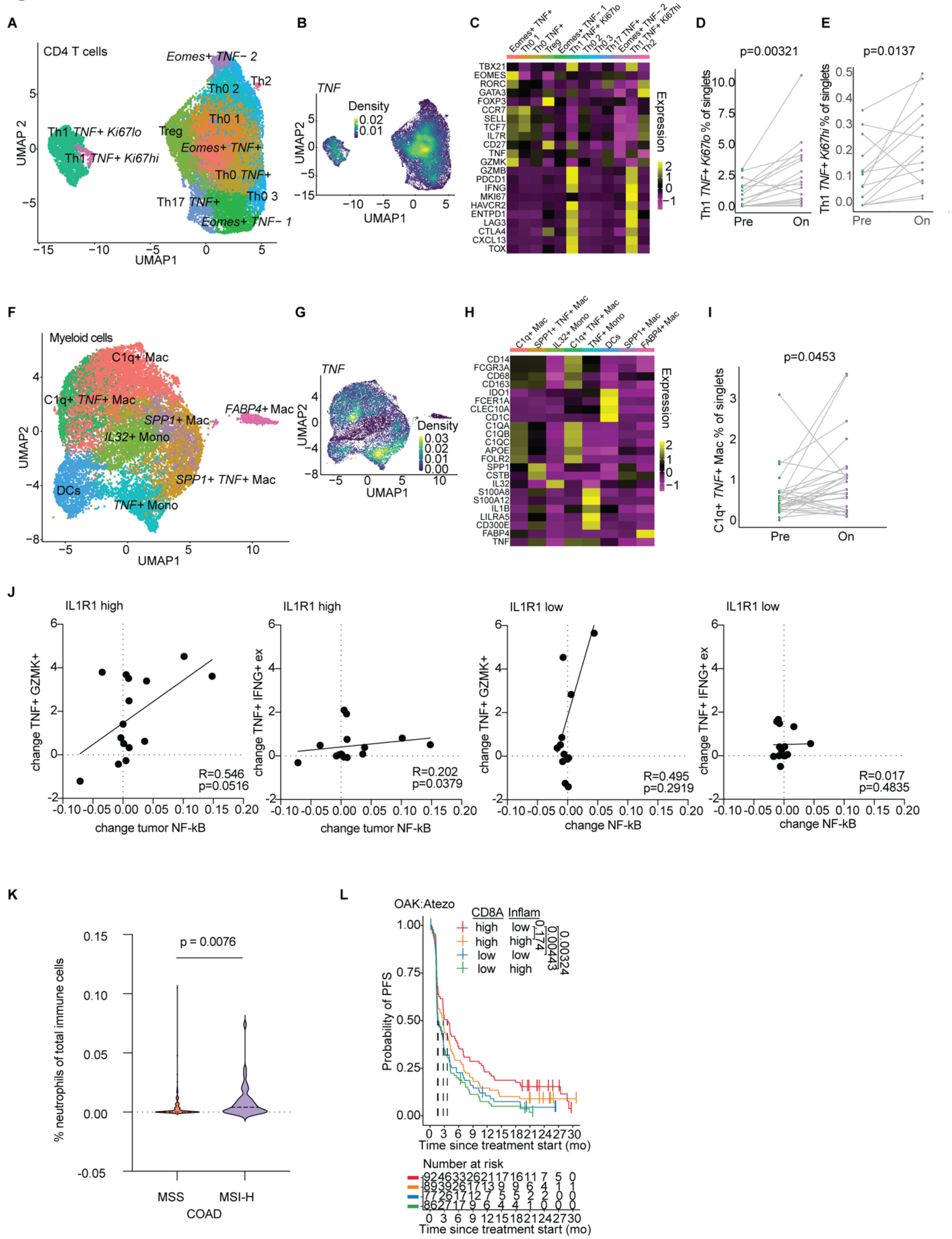


140

**Supplementary Figure S11.**

**A,** UMAP of 24,468 CD4 T cells from the BIOKEY study (EGAS00001004809), color coded by CD4 T cell phenotype. **B,** UMAP cell density plot for *TNF* expressing cells. **C,** Heatmap of genes defining CD4 T cell clusters. **D and E,** Percent of cells in indicated cluster out of singlet cells in tumor for pre vs. on-treatment samples for each patient. P values by two-tailed matched-pairs t test. **F,** UMAP of 16,485 myeloid cells from the BIOKEY study (EGAS00001004809), color coded by myeloid cell phenotype. **G,** UMAP cell density plot for *TNF* expressing myeloid cells. **H,** Heatmap for genes defining myeloid cell clusters. **I,** Percent of cells in the C1q+ *TNF*+ Macrophage cluster out of singlet cells in tumor for pre vs. on-treatment samples for each patient. P value by two-tailed matched-pairs t test. **J,** Spearman correlation analysis between the changes in indicated cluster frequencies vs. changes in tumor NF-kB score before and after ICI treatment, stratified by *IL1R1* high and *IL1R1* low status. **K,** Violin plots of neutrophil quantification by CIBERTSORTx from MSS and MSI-H COAD patients in TCGA. P value by two-tailed Mann-Whitney test. **L,** Kaplan-Meier curves of patients in the OAK trial (NCT02008227) for progression-free survival (PFS), stratified by *CD8A* status and inflammation gene score status (“inflam”). P values by log-rank test.

#### Figure S12

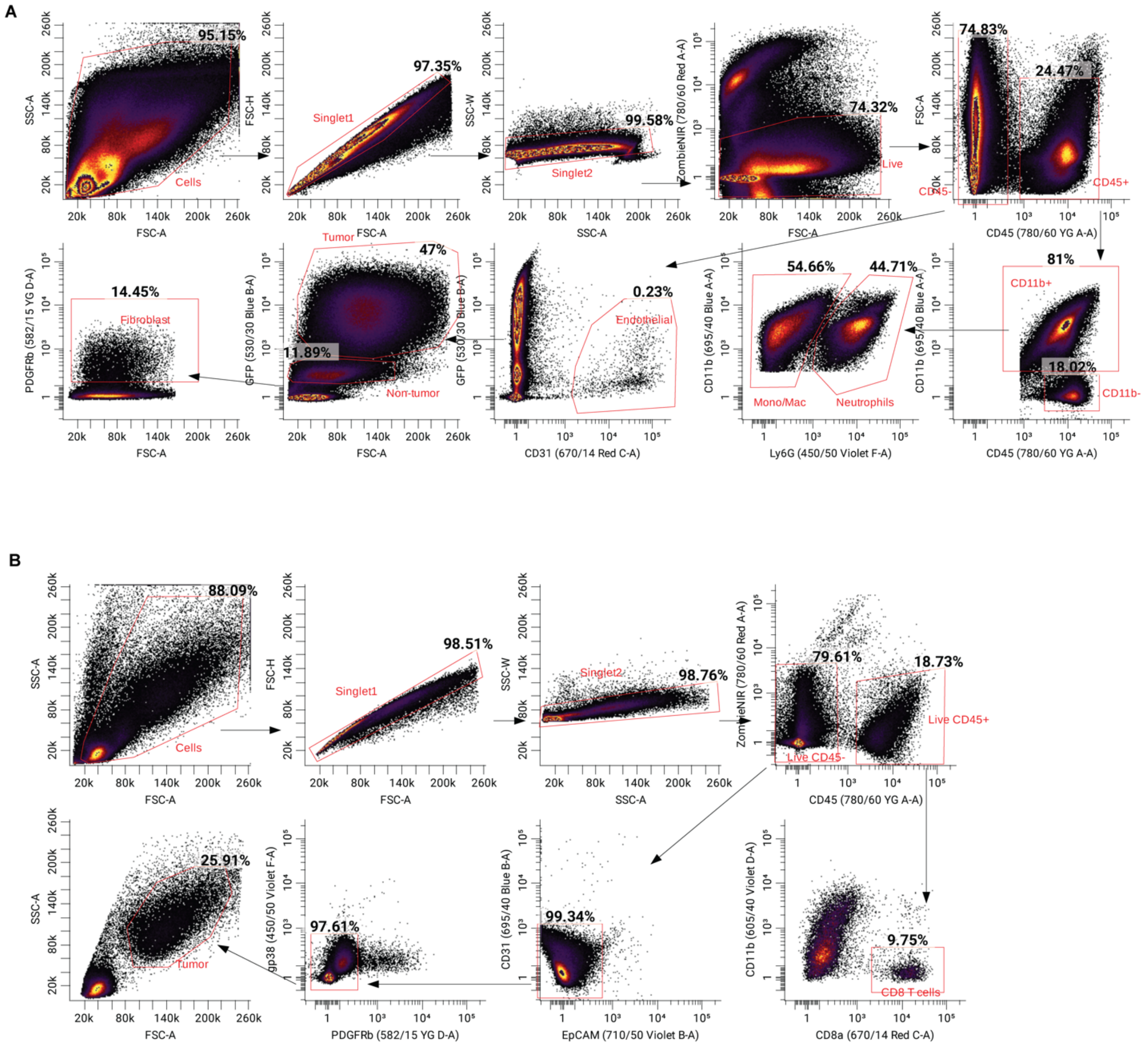

A
95.15%
Cells
SSC-A
FSC-A
97.35%
Singlet1
FSC-H
99.58%
Singlet2
SSC-W
SSC-A
74.32%
Live
ZombieNIR (780/60 Red A-A)
74.83%
24.47%
CD45-
CD45+
CD45 (780/60 YG A-A)
81%
CD11b+
18.02%
CD11b-
CD11b (695/40 Blue A-A)
54.66%
44.71%
Mono/Mac
Neutrophils
CD11b (695/40 Blue A-A)
Ly6G (450/50 Violet F-A)
0.23%
Endothelial
GFP (530/30 Blue B-A)
CD31 (670/14 Red C-A)
47%
Tumor
11.89%
Non-tumor
14.45%
Fibroblast
PDGFRb (582/15 YG D-A)
B
88.09%
98.51%
98.76%
79.61%
18.73%
Live CD45-
Live CD45+
9.75%
CD8 T cells
CD11b (605/40 Violet D-A)
CD8a (670/14 Red C-A)
99.34%
CD31 (695/40 Blue B-A)
EpCAM (710/50 Violet B-A)
97.61%
gp38 (450/50 Violet F-A)
PDGFRb (582/15 YG D-A)
25.91%

**Supplementary Figure S12**

**A,** Gating strategy used for experiment in **Fig. 3F**. **B,** Gating strategy used for CD8 T cell and tumor co-culture experiments shown in **Fig. 5D**.

#### Figure S13

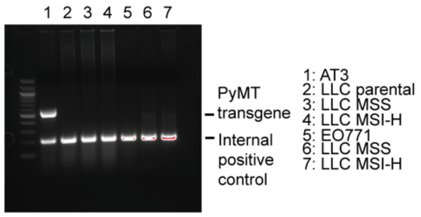


160

**Supplementary Figure S13.**

Gel electrophoresis of PCR for the PYMT transgene (top band) and internal control (bottom bands) from indicated cell lines.

**Supplementary Table S1. CRISPR-Cas9 reagents and parameters**

| | |
|---|---|
| Scrambled nontargeting control | GGTTCTTGACTACCGTAATT |
| LLC MSH2 crRNA combination | GGTTAATACCCTGATACAGT + CACATGAGCGAAGCTAACAA |
| B16F10 MSH2 crRNA combination | CCTTAATAAATGCAGCCCGG + GGTTAATACCCTGATACAGT |
| IL1R1 crRNA combination | AGCATACAATTGTAGCCGTG + CAGCAAGACCCCCATATCAG |
| ssODN | TTAGCTCTGTTTACGTCCCAGCGGGCATGAGAGTAACAAGAG GGTGTGGTAATATTACGGTACCGAGCACTATCGATACAATAT GTGTCATACGGACACG |
| LLC buffer and code | Buffer SE, code DS-120 |
| B16F10 buffer and code | Buffer SG, code DC-135 |

**Supplementary Table S2. PCR primer sequences**

| | |
|---|---|
| AA003063 F | NED-ACGTCAAAAATCAATGTTAGG |
| AA003063 R | CAGCAAGGGTCCCTGTCTTA |
| AC096777 F | VIC- TCCCTGTATAACCCTGGCTGACT |
| AC096777 R | GCAACCAGTTGTCCTGGCGTGGA |
| L24372 F | FAM-GGGAAGACTGCTTAGGGAAGA |
| L24372 R | ATTTGGCTTTCAAGCATCCATA |
| U12235 F | VIC-GCTCATCTTCGTTCCCTGTC |
| U12235 R | CATTCGGTGGAAAGCTCTGA |
| PYMT transgene forward | GGAAGCAAGTACTTCACAAGGG |
| PYMT transgene reverse | GGAAAGTCACTAGGAGCAGGG |
| Internal positive control forward | CAAATGTTGCTTGTCTGGTG |
| Internal positive control reverse | GTCAGTCGAGTGCACAGTTT |

**Supplementary Table S3. Table of cytokines analyzed in Fig. 3D**

| Cytokine (pg/mL) | IgG | | | | | ICIs | | | | | |
|---|---|---|---|---|---|---|---|---|---|---|---|
| 6Ckine/Exodus 2 | 304.5 | 669.86 | 305.25 | 309.24 | 984.62 | 118.72 | 1163.68 | 815.55 | 1276.75 | 166.26 | 176.97 |
| Eotaxin | 17.65 | 48.93 | 27.53 | 45.21 | 60.82 | 22.48 | 65.94 | 53.92 | 79.99 | 45.89 | 21.97 |
| Fractalkine | 17.61 | 13.76 | 0 | 8.85 | 13.76 | 2.54 | 18.86 | 15.07 | 12.41 | 15.71 | 15.07 |
| G-CSF | 16.18 | 72.28 | 52.08 | 11.73 | 159.72 | 108.55 | 553.77 | 539.23 | 29.03 | 183.54 | 843.93 |
| GM-CSF | 7.71 | 17.3 | 9.38 | 9.38 | 0 | 25.52 | 54.56 | 62.05 | 9.38 | 44.5 | 24.88 |
| IFNb-1 | 29.58 | 61.34 | 52.71 | 38.27 | 39.72 | 38.27 | 69.96 | 67.09 | 46.94 | 52.71 | 36.82 |
| IFNg | 1.84 | 3.22 | 1.11 | 4.15 | 0 | 0 | 5.19 | 8.89 | 0 | 3.75 | 5.64 |
| IL-10 | 4.18 | 6.08 | 5.24 | 8.53 | 2.65 | 5.24 | 10.13 | 4.82 | 9.13 | 7.31 | 5.24 |
| IL-11 | 22.28 | 27.8 | 25.5 | 28.72 | 19.99 | 22.28 | 156.45 | 94.1 | 58.34 | 59.27 | 68.62 |
| IL-12p40 | 4.49 | 11.01 | 6.84 | 4.49 | 2.89 | 0 | 3.5 | 2.07 | 8.18 | 5.28 | 3.5 |
| IL-12p70 | 0 | 0.58 | 0 | 0 | 2.54 | 0 | 3.2 | 0 | 1.89 | 8.42 | 0 |
| IL-13 | 0.47 | 0 | 0 | 0 | 0.62 | 0 | 0 | 0.03 | 0.28 | 0 | 0.47 |
| IL-15 | 19.31 | 27.37 | 22.57 | 20.94 | 35.18 | 28.95 | 25.78 | 22.57 | 35.18 | 28.95 | 12.64 |
| IL-16 | 437.62 | 505.73 | 370.53 | 422.44 | 412.24 | 413.88 | 482.31 | 471.85 | 390.92 | 411.52 | 529.22 |
| IL-17 | 0 | 0 | 0 | 0 | 0 | 0 | 0 | 0 | 0 | 0 | 0 |
| IL-1a | 0 | 0 | 0 | 0 | 0 | 0 | 0 | 0 | 87.4 | 0 | 0 |
| IL-1b | 4.82 | 5.32 | 5.48 | 4.15 | 2.39 | 6.77 | 16.02 | 9.57 | 8.19 | 7.09 | 6.13 |
| IL-2 | 16.1 | 13.03 | 9.86 | 16 | 13.15 | 17.68 | 17.27 | 12.92 | 20.23 | 18.81 | 13.6 |
| IL-3 | 0 | 0.07 | 0 | 0.11 | 0.26 | 0.72 | 0.22 | 0.41 | 0 | 0.41 | 0 |
| IL-4 | 0.53 | 0.67 | 0.25 | 0.06 | 0.25 | 0.39 | 0.56 | 0.45 | 0.64 | 0.2 | 0 |
| IL-5 | 1.05 | 1.7 | 1.34 | 1.48 | 0.89 | 1.05 | 1.95 | 1.15 | 1.1 | 1.74 | 1.05 |
| IL-6 | 6.27 | 20.27 | 6.77 | 3.99 | 8.08 | 29.24 | 52.89 | 60.43 | 4.72 | 30.32 | 21.73 |
| IL-7 | 4.96 | 4.13 | 2.28 | 2.28 | 3.62 | 4.96 | 7.96 | 3.96 | 8.8 | 5.63 | 2.28 |
| IL-9 | 29.51 | 71.77 | 51.28 | 62.17 | 24.62 | 60.35 | 35.33 | 18.79 | 58.52 | 50.96 | 48.68 |
| IP-10 | 152.33 | 406.81 | 82.38 | 170.22 | 99.18 | 225.14 | 220.61 | 160.16 | 234.88 | 259.51 | 222.16 |
| CXCL1 | 47.35 | 194.79 | 130.5 | 39.57 | 72.74 | 258.29 | 650.34 | 399.14 | 56.96 | 398.71 | 404.64 |
| LIF | 9.46 | 8.86 | 16.95 | 5.95 | 14.6 | 13.65 | 53.9 | 34.79 | 11.33 | 20.04 | 48.77 |
| LIX | 0 | 0 | 0 | 0 | 0 | 0 | 0 | 0 | 0 | 0 | 0 |
| M-CSF | 10.45 | 21.22 | 14.38 | 7.66 | 9.46 | 16.72 | 17.88 | 24.09 | 11.38 | 17.36 | 16.07 |
| MCP-1 | 548.89 | 964.97 | 702.81 | 641.77 | 757.97 | 954.14 | 1552.56 | 289.38 | 612.76 | 1243.18 | 1379.36 |
| MCP-5 | 976.58 | 1797.57 | 751.15 | 572.35 | 1055.91 | 725.85 | 1435.13 | 1018.87 | 1269.19 | 1499.34 | 712 |
| MDC | 5.72 | 5.56 | 4.08 | 5.08 | 4.86 | 4.86 | 6.29 | 5.07 | 4.25 | 6.59 | 2.8 |
| MIG | 809.39 | 552.49 | 255.45 | 805.26 | 154.66 | 1082.98 | 1585.34 | 1166.23 | 585.38 | 922 | 949.54 |
| MIP-1a | 18.18 | 25.78 | 26.22 | 16.97 | 16.97 | 24.41 | 36.19 | 34.1 | 27.94 | 38.19 | 34.1 |
| MIP-1b | 3.6 | 8.87 | 1.15 | 0 | 0 | 5.04 | 18.47 | 0 | 0 | 11.6 | 5.04 |
| MIP-2 | 273.85 | 0 | 35.15 | 128.88 | 551.65 | 305.06 | 1734.96 | 1903.9 | 0 | 839.87 | 721.32 |
| MIP-3a | 1 | 1.66 | 1.66 | 1.33 | 1.61 | 1.33 | 2.11 | 1.58 | 2.42 | 2.06 | 1.36 |
| MIP-3b | 27.74 | 97.17 | 51.46 | 26.75 | 28.97 | 64.2 | 61.09 | 88.14 | 34.49 | 69.58 | 31.69 |

| | | | | | | | | | | | |
|---|---|---|---|---|---|---|---|---|---|---|---|
| RANTES | 1.83 | 1.35 | 1.76 | 1.9 | 1.47 | 1.27 | 3.28 | 1.93 | 2.43 | 2.43 | 2.06 |
| TARC | 40.06 | 26.49 | 18.76 | 28.19 | 22.37 | 27.99 | 26.68 | 27.83 | 21.78 | 32.47 | 16.59 |
| TIMP-1 | 904.9 | 1599.77 | 1154.13 | 1146.85 | 1177.39 | 1202.23 | 1389.59 | 1512.61 | 1910.66 | 1268.95 | 1266.35 |
| TNFa | 1.98 | 1.81 | 1.16 | 1.67 | 1.47 | 1.9 | 2.86 | 1.71 | 3.25 | 3.56 | 1.71 |
| VEGF | 2.67 | 3.62 | 6.98 | 1.78 | 5.42 | 3.77 | 19.32 | 8.72 | 4.08 | 7.79 | 9.07 |

172

**Supplementary Table S4. Mutation analysis of cell line exome sequencing data.**

| Mutation | #CHROM | POS | REF | ALT | 4T1 | B16F10 | CT26 | LLC | LLC MSS |
|---|---|---|---|---|---|---|---|---|---|
| Kras G12C | 6 | 145192498 | C | A | 0/0 | 0/0 | 0/0 | **0/1** | **0/1** |
| EGFR P992P | 11 | 16859755 | A | G | 0/0 | 0/0 | 0/0 | **0/1** | **0/1** |

**Supplementary Table S5. CyTOF antibody panel**

| Mass | Metal | Protein | Titrated Conc. (ug/ml) | Clone | Isotype | Catalog | Vendor | RRID |
|---|---|---|---|---|---|---|---|---|
| 89 | Y | CD45 | 1.5 | 30-F11 | rat IgG2b | 103102 | Biolegend | AB_312967 |
| 111 | Cd | CD3 | 6 | 17A2 | rat IgG2b | 100202 | Biolegend | AB_312659 |
| 112 | Cd | CD64 | 6 | X54--5/7.1 | mouse IgG1 | 139302 | Biolegend | AB_10613107 |
| 113 | In | Ter119 | 3 | TER-119 | Rat IgG2b | 116202 | Biolegend | AB_313703 |
| 114 | Cd | CD8 | 3 | 53-6.7 | rat IgG2a | 100702 | Biolegend | AB_312741 |
| 116 | Cd | CD4 | 1.5 | RM-5 | rat IgG2a | 100506 | Biolegend | AB_312709 |
| 139 | La | Ly6G | 0.375 | 1A8 | rat IgG2a | 127626 | Biolegend | AB_2561340 |
| 140 | Ce | KLRG1 | 1.5 | 2F1 | syrian hamster IgG2 | 562190 | BD biosciences | AB_11154418 |
| 141 | Pr | granzyme B | 3 | QA16A02 | mouse IgG1 | 372202 | Biolegend | AB_2686929 |
| 142 | Nd | CD49b | 0.75 | HMa2 | armenian hamster IgG | 103501 | Biolegend | AB_313024 |
| 143 | Nd | CD11c | 0.75 | N418 | armenian hamster IgG | 117341 | Biolegend | AB_2562807 |
| 144 | Nd | CD206 | 0.75 | C068C2 | rat IgG2a | 141702 | Biolegend | AB_10900233 |
| 145 | Nd | CD27 | 6 | LG 3A10 | armenian hamster IgG | 124202 | Biolegend | AB_1236456 |
| 146 | Nd | CD138 | 6 | 281-2 | rat IgG2a | 142502 | Biolegend | AB_10965646 |
| 147 | Sm | PD-L1 | 3 | 10F.9G2 | rat IgG2b | 124302 | Biolegend | AB_961228 |
| 148 | Nd | ICOS | 3 | C398.4A | armenian hamster IgG | 313502 | Biolegend | AB_416326 |
| 149 | Sm | SiglecF | 1.5 | E50-2440 | rat IgG2a | 552125 | BD biosciences | AB_394340 |
| 150 | Nd | PDCA-1 | 0.75 | 120G8.04 | rat IgG1 | DDX0390P-100 | Novus Biologicals | AB_2827525 |
| 151 | Eu | Ly6C | 0.375 | HK1.4 | rat IgG2c | 128002 | Biolegend | AB_1134214 |
| 152 | Sm | Ki67 | 1.5 | SolA15 | rat IgG2a | 14-5698-82 | Thermo Fisher | AB_10854564 |
| 153 | Eu | CD11b | 1.5 | HK1.4 | rat IgG2b | 101202 | Biolegend | AB_312785 |
| 154 | Sm | cKit | 1.5 | 2B8 | rat IgG2b | 105802 | Biolegend | AB_313211 |
| 155 | Gd | CD103 | 1.5 | 2E7 | armenian hamster IgG | 121402 | Biolegend | AB_535945 |
| 156 | Gd | CD80 | 0.1875 | 16-10A1 | armenian hamster IgG | 104702 | Biolegend | AB_313123 |
| 157 | Gd | CD83 | 1.5 | Michel-19 | rat IgG1 | 121502 | Biolegend | AB_572023 |
| 158 | Gd | B220 | 1.5 | RA3-6B2 | rat IgG2a | 103202 | Biolegend | AB_312987 |
| 159 | Tb | PD-1 | 0.375 | 29F.1A12 | rat IgG2a | 135202 | Biolegend | AB_1877121 |
| 160 | Gd | NK1.1 | 1.5 | PK136 | mouse IgG2a | 108702 | Biolegend | AB_313389 |
| 161 | Dy | T-bet | 6 | 4B10 | mouse IgG1 | 644825 | Biolegend | AB_2563788 |

| | | | | | | | | |
|---|---|---|---|---|---|---|---|---|
| 162 | Dy | TCRgd | 0.75 | GL3 | armenian hamster IgG | 118101 | Biolegend | AB_313826 |
| 163 | Dy | CD62L | 6 | 95218 | rat IgG2a | MAB5761 | R&D Systems | AB_2254524 |
| 164 | Dy | TCF1 | 1.5 | 2203 | rabbit IgG | 2203S | Cell Signaling | AB_2199302 |
| 165 | Ho | CD69 | 3 | H1.2F3 | armenian hamster IgG | 104502 | Biolegend | AB_313105 |
| 166 | Er | FcER1a | 3 | MAR-1 | armenian hamster IgG | 134321 | Biolegend | AB_2563768 |
| 167 | Er | Foxp3 | 3 | NRRF-30 | rat IgG2a | 14-4771-80 | Thermo Fisher | AB_529583 |
| 168 | Er | CD86 | 0.1875 | GL-1 | rat IgG2a | 105002 | Biolegend | AB_313145 |
| 169 | Tm | F4/80 | 6 | BM8 | rat IgG2a | 123143 | Biolegend | AB_2563767 |
| 170 | Er | CD115 | 1.5 | AFS98 | rat IgG2a | 135521 | Biolegend | AB_2563709 |
| 171 | Yb | CD38 | 0.75 | 90 | rat IgG2a | 102723 | Biolegend | AB_2563746 |
| 172 | Yb | TIM3 | 3 | RMT3-23 | rat IgG2a | 119702 | Biolegend | AB_345376 |
| 173 | Yb | CD19 | 0.375 | 6D5 | rat IgG2a | 115547 | Biolegend | AB_2562806 |
| 174 | Yb | iNOS | 0.75 | CXNFT | rat IgG2a | 14-5920-82 | Thermo Fisher | AB_2572890 |
| 175 | Lu | CD44 | 0.375 | IM7 | rat IgG2b | 103002 | Biolegend | AB_312953 |
| 176 | Yb | CD39 | 0.75 | 24DMS1 | rat IgG2b | 14-0391-82 | Thermo Fisher | AB_1210501 |
| 209 | Bi | MHC II | 0.1 | M5/114.15.2 | rat IgG2b | 107602 | Biolegend | AB_313317 |

**Supplementary Table S6. Flow cytometry antibodies**

| Target | Fluorophore | Vendor | Clone | Catalog | RRID |
|---|---|---|---|---|---|
| BrdU | APC | Thermo Fisher | | 8817-6600-42 | AB_2575274 |
| CD11b | BV711 | Biolegend | M1/70 | 101241 | AB_2563310 |
| CD11b | PerCP-Cy5.5 | Biolegend | M1/70 | 101228 | AB_893232 |
| CD31 | APC | Biolegend | 390 | 102410 | AB_312905 |
| CD31 | PerCP-Cy5.5 | Biolegend | 390 | 102420 | AB_10613644 |
| CD4 | BV510 | Biolegend | RM4-5 | 100559 | AB_2562608 |
| CD45 | PE-Cy7 | Biolegend | 30-F11 | 103114 | AB_312979 |
| CD45.1 | PE | Biolegend | A20 | 110708 | AB_313497 |
| CD8a | APC | Biolegend | 53-6.7 | 100712 | AB_312751 |
| CD8a | BV650 | Biolegend | 53-6.7 | 100741 | AB_2563056 |
| CD8a | PerCP-Cy5.5 | Biolegend | 53-6.7 | 100734 | AB_2075238 |
| EpCAM | BV605 | Biolegend | G8.8 | 118227 | AB_2563984 |
| F4/80 | PerCP-Cy5.5 | Biolegend | BM8 | 123127 | AB_893484 |
| gp38 | BV421 | Biolegend | 8.1.1 | 127423 | AB_2814017 |
| H-2Kb/H-2Db | FITC | Biolegend | 28-8-6 | 114605 | AB_313597 |
| IkBa | AF488 | Cell Signaling Technology | 93H1 | 4886S | AB_390789 |
| Ly6G | BV421 | Biolegend | 1A8 | 127627 | AB_2562567 |
| Ly6G | PE-Cy7 | Biolegend | 1A8 | 127618 | AB_1877261 |
| NK1.1 | PE | BD Biosciences | PK136 | 553165 | AB_394677 |
| PDGFRb | PE | Biolegend | APB5 | 136006 | AB_1953271 |
| TCRb | BV421 | Biolegend | H57-597 | 109230 | AB_2562562 |
| TNFα | APC | Biolegend | MP6-XT22 | 506308 | AB_315429 |
| TNFα | FITC | Biolegend | MP6-XT22 | 506304 | AB_315425 |
